# Mandatory coupling of zygotic transcription to DNA replication in early *Drosophila* embryos

**DOI:** 10.1101/2022.01.04.474810

**Authors:** Chun-Yi Cho, James P. Kemp, Robert J. Duronio, Patrick H. O’Farrell

## Abstract

Collisions between transcribing RNA polymerases and DNA replication forks are disruptive. The threat of collisions is particularly acute during the rapid early embryonic cell cycles of *Drosophila* when S phase occupies the entirety of interphase. We hypothesized that collision-avoidance mechanisms safeguard the onset of zygotic transcription in these cycles. To explore this hypothesis, we used real-time imaging of transcriptional events at the onset of each interphase. Endogenously tagged RNA polymerase II (RNAPII) abruptly formed clusters before nascent transcripts accumulated, indicating recruitment prior to transcriptional engagement. Injection of inhibitors of DNA replication prevented RNAPII clustering, blocked formation of foci of the pioneer factor Zelda, and largely prevented expression of transcription reporters. Knockdown of Zelda or the histone acetyltransferase CBP prevented RNAPII cluster formation except at the replication-dependent (RD) histone gene locus. We suggest a model in which the passage of replication forks allows Zelda and a distinct pathway at the RD histone locus to reconfigure chromatin to nucleate RNAPII clustering and promote transcriptional initiation. The replication dependency of these events defers initiation of transcription and ensures that RNA polymerases transcribe behind advancing replication forks. The resulting coordination of transcription and replication explains how early embryos circumvent collisions and promote genome stability.

## INTRODUCTION

Reading of the genetic code by both replicative and transcriptional machineries can create conflicts, yet the mechanisms that coordinate template use remain incompletely understood. The expected incompatibility of simultaneous use of the template for replication and transcription was first demonstrated in vitro in a T4 bacteriophage system nearly 40 years ago (Bedinger et al., 1983). Collisions, especially head-on collisions between advancing RNA polymerases and replication forks, disrupt both transcription and replication. Such collisions are now known to occur in both prokaryotes and eukaryotes. Bacteria have evolved to co-orient the majority of genes with the direction of replication on their circular genomes, thus minimizing head-on collisions (Merrikh et al., 2012). Cells have also evolved numerous mechanisms to resolve collisions when they do occur and to mitigate the genome instability that can result (García-Muse and Aguilera, 2016). Yet beyond this, we know little about how replication and transcription might be coordinated to minimize disruptive conflicts.

Coordinating DNA replication and transcription is especially critical yet challenging during the early development of multicellular organisms. In many species, including *Drosophila*, the embryos undergo a series of rapid cleavage cycles after fertilization. During these cycles, nuclei alternate between S phase and mitosis without gap phases. Although transcription is minimal in the first few cycles, a process called zygotic genome activation (ZGA) leads to a minor wave of gene expression before the cell cycle significantly slows down (Schulz and Harrison, 2019). For example, zygotic transcription in young *Drosophila* embryos can be detected as early as during nuclear cycle (nc) 8 (ten Bosch et al., 2006; Erickson and Cline, 1993; Kwasnieski et al., 2019), when the interphase is about 4 minutes and is occupied entirely by an S phase that begins with nearly simultaneous activation of origins throughout the genome (Blumenthal et al., 1974; Ji et al., 2004). As the minor wave of zygotic transcription lays the foundation for embryogenesis, its success is crucial. We sought to understand how it is coordinated with DNA replication during nuclear cycles 10-13.

The *Drosophila* embryo provides an ideal system to probe for mechanisms that mitigate conflicts between replication and transcription. Previous studies have separately examined the real-time dynamics of DNA replication and transcription in the syncytial-blastoderm embryos of *Drosophila* going through nuclear cycles (Garcia et al., 2013; Lucas et al., 2013; McCleland et al., 2009; Shermoen et al., 2010). The data support a temporal order where DNA replication begins immediately during mitotic exit, while active transcription occurs a few minutes later. Furthermore, ATAC-seq data from staged embryos suggest that chromatin accessibility declines in early interphase and is restored with a delay (Blythe and Wieschaus, 2016). Orderly nucleosome positioning around enhancers and promoters is disrupted upon passage of replication forks, and the restoration of order can be slow (Ramachandran and Henikoff, 2016). Several studies have suggested that a class of transcription factors called pioneer factors are specialized for restoring a chromatin configuration that is permissive for regulated transcription (Blythe and Wieschaus, 2016; McDaniel et al., 2019; Sun et al., 2015; Zaret and Carroll, 2011). Nonetheless, the exact interplay between DNA replication, transcription, and pioneer factors in early embryos remains unknown.

Several transcription factors have been implicated in pioneering ZGA in *Drosophila* (Duan et al., 2021; Gaskill et al., 2021; Liang et al., 2008). Among them, the zinc-finger transcription factor Zelda (Zld) has been the most extensively studied. The mRNA encoding Zld is supplied maternally and expressed ubiquitously throughout the early embryo (Liang et al., 2008; Nien et al., 2011). Zld protein binds to thousands of enhancers and promoters during minor ZGA, and its binding sites exhibit increased chromatin accessibility and histone acetylation (Harrison et al., 2011; Li et al., 2014; Nien et al., 2011; Schulz et al., 2015; Sun et al., 2015). The depletion of maternally expressed Zelda significantly reduces the level of zygotic transcription, and the embryos become highly defective at the mid-blastula transition (MBT) and fail at cellularization and gastrulation (Liang et al., 2008). More recently, live microscopy revealed that fluorescently tagged Zld forms dynamic hubs in the nucleus (Dufourt et al., 2018; Mir et al., 2018). In addition, Zld can increase the local concentration of other transcription factors and facilitate their binding to target DNA (Mir et al., 2017, 2018; Yamada et al., 2019). These observations are in line with recent exciting findings from mammalian cells, which suggest a model that transcription occurs within nuclear microenvironments formed through liquid-liquid phase separation (Boehning et al., 2018; Boija et al., 2018; Cho et al., 2018; Sabari et al., 2018). The formation of such transcriptional condensates is facilitated by the intrinsically disordered regions (IDRs) present in many transcription factors and co-activators, as well as by the carboxy-terminal domain (CTD) of RNA polymerase II (RNAPII) (Cramer, 2019; Hnisz et al., 2017). However, it remains unclear how the formation of transcriptional nuclear microenvironments is coordinated with cell-cycle events, and the dynamics of their establishment have yet to be fully elucidated.

In this study, we investigate events contributing to transcriptional initiation and determine their dependency on replication in early *Drosophila* embryos. We show that endogenously tagged RNAPII forms cell-cycle-coordinated clusters during ZGA. The formation of RNAPII clusters precedes the onset of transcription in each cycle and depends on multiple regulators, including the pioneer factor Zld. Inhibiting DNA replication blocks the clustering of both Zld and RNAPII and prevents the onset of transcription following mitotic exit. Our data suggest that Zld-dependent clustering of RNAPII promotes a “burst” of transcriptional initiation during the short interphases, and that DNA replication is required to initiate this process. We propose that embryonic cells largely avoid transcription-replication conflicts by enforcing a temporal order in which newly initiated replication forks are followed by transcribing RNA polymerases moving at comparable speeds, thereby reducing collisions.

## RESULTS

### RNA polymerase II (RNAPII) forms two distinct classes of clusters in blastoderm-stage *Drosophila* embryos

Motivated by recent reports on the clustering or phase separation of RNA polymerase II (RNAPII) during transcription in mammalian cell lines (Boehning et al., 2018; Cho et al., 2016, 2018; Cisse et al., 2013), we used CRISPR/Cas9 to tag the N-terminal end of two RNAPII subunits, Rpb1 and Rpb3, with mCherry or EGFP, respectively, for live imaging of RNAPII localization during the early embryonic nuclear cycles of *Drosophila*. Despite a slightly lower hatch rate of embryos laid by homozygous EGFP-Rpb3 females, flies homozygous for either mCherry-Rpb1 or EGFP-Rpb3 are healthy and fertile (Figure S1A). Therefore, the two tagged RNAPII subunits largely retain their original function.

Embryos from mothers expressing mCherry-Rpb1 or EGFP-Rpb3 exhibited nuclear foci of these RNAPII subunits during syncytial-blastoderm nuclear cycles (e.g. Figure 1A). In embryos co-expressing the two markers, the nuclear foci of Rpb1 and Rpb3 colocalized (Figure 1B), indicating that foci represent clusters of RNAPII holoenzymes rather than aggregates of fluorescent proteins. We detected sparse nuclear foci as early as nc10 when nuclei migrated from the interior of the embryo to the surface around the beginning of zygotic genome activation (Figure S1B). Live imaging revealed the dynamics of RNAPII clustering. Two distinct classes of clusters with different cell-cycle behaviors were apparent (Figure 1A and Movie S1). During the first minute following each mitosis, RNAPII accumulated uniformly in the newly formed nucleus. Between 1-3 minutes into interphase, small RNAPII clusters began to appear throughout the nucleus. Their numbers and intensity peaked between 3-4 minutes. Most of the small clusters then dissolved and largely disappeared by early prophase (Figure 1A, see 7-min frame of nc11 for example). One or two clusters in each nucleus were distinctive—they increased in size throughout interphase, were much larger than the dispersed clusters, persisted through prophase, and declined in prometaphase (Figure S1C and Movie S1).

**Figure 1.**
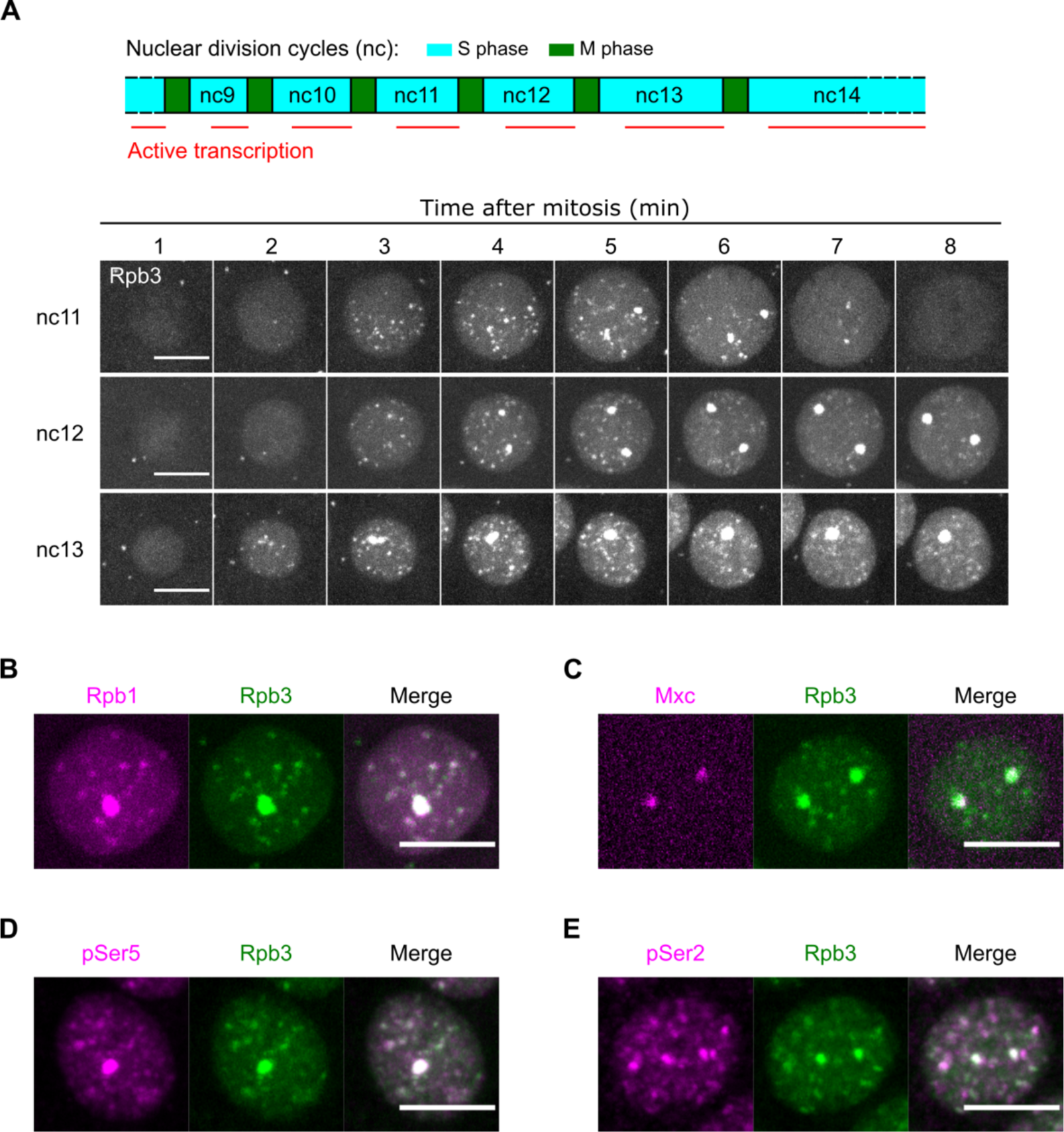
Two classes of RNAPII clusters appear during zygotic genome activation in *Drosophila* embryos. (A, top) Schematic illustrating the temporal overlap between nuclear division cycles and the onset of zygotic transcription in early *Drosophila* embryos. (A, bottom) Representative stills from live imaging of endogenously tagged EGFP-Rpb3 during nuclear cycles (nc) 11-13. Maximal projections across the entire nuclei are shown. In each cycle, images centering on a single nucleus are cropped and displayed. The 0-minute timepoint is determined manually using His2Av-mRFP as shown in Figure S1C. (B-C) Confocal live imaging of EGFP-Rpb3 embryos co-expressing either mCherry-Rpb1 or Mxc-mScarlet during nuclear division cycles. (D-E) Immunostaining of the EGFP-Rpb3 embryos using an anti-GFP antibody with another antibody recognizing either the phospho-Ser5 or phosphor-Ser2 modifications in the CTD repeats of Rpb1. All scale bars in Figure 1 indicate 5 μm. See also Figures S1, S2, and Movie S1.

The progressive maturation of the large RNAPII clusters from nc11 to nc13 resembled that of histone locus bodies (HLBs), which is a nuclear body that regulates histone biogenesis (Duronio and Marzluff, 2017; Hur et al., 2020; Nizami et al., 2010). In *Drosophila*, clusters of RD histone genes are organized in a tandem array of about 100 copies at the *HisC* locus. Onset of histone gene transcription during ZGA coincides with the recruitment of HLB components to the *HisC* locus (Edgar and Schubiger, 1986; White et al., 2011). We found that the two large EGFP-Rpb3 foci colocalized with mScarlet-tagged Mxc, an essential scaffolding protein of HLBs (Figure 1C)(Huang et al., 2021; Kemp et al., 2021; Terzo et al., 2015). Thus, the large RNAPII clusters mark the HLBs, and the components of HLBs might modulate the distinct cell-cycle dynamics of RNAPII at the RD histone locus (Hur et al., 2020).

We next asked whether both large and small RNAPII clusters contained active forms of RNAPII. During the transcriptional cycle, the CTD domain of the Rpb1 subunit undergoes different post-translational modifications. Among them, phosphorylation of serine 5 (pSer5) sites in the CTD heptapeptide repeats is enriched near promoter-proximal regions during initiation and stalling, and phosphorylation of serine 2 (pSer2) sites in the CTD increases during transcriptional elongation across the gene body (Harlen and Churchman, 2017). Using immunostaining, we showed that all large RNAPII clusters at HLBs also stained for pSer5 and pSer2 (Figures 1D and 1E)(Kemp et al., 2021). While the small RNAPII clusters showed frequent overlap with pSer5 and pSer2 staining (Figures 1D and 1E), the relative intensity of co-staining varied between foci particularly for pSer2 (Figure S2). The rapid emergence and disappearance of individual RNAPII foci (Figure 1A) suggest that this heterogeneity in staining represents asynchronous maturation of different foci. Most strong EGFP-Rpb3 signals have associated pSer5 signal; however, the associated pSer5 signal varies from weak to strong, consistent with initial recruitment of unmodified RNAPII followed promptly by progressive Ser5 phosphorylation (Forero-Quintero et al., 2021). The alignment of EGFP-Rpb3 and pSer2 signals differs in two ways. First, there are frequent EGFP-Rpb3 foci with weak pSer2 signals. This staining pattern suggests recruitment of some RNAPII without concomitant transcription. Second, some of the brightest pSer2 staining foci have weak to very weak EGFP-Rpb3 signal, suggesting transcription with little evident recruited polymerase. This latter observation suggests that much of the RNAPII recruited to clusters might disperse with only a minority of the polymerases progressing to full Ser2 phosphorylation of CTD repeats. We conclude that the cell-cycle-coordinated clustering of nuclear RNAPII accompanies minor ZGA in *Drosophila* and suggest that cluster formation is followed by a dynamic cascade of modification independently occurring at different loci.

### RNAPII clustering precedes transcriptional engagement

Classically, RNAPII binds at promoters and initiates transcription. Accordingly, one RNAPII would be recruited to the gene with each initiation, and promoter recruitment of the subsequent RNAPII would be delayed at least until the previously engaged polymerase had moved out of the way. This model predicts that polymerase recruitment would be progressive and occur in parallel to the accumulation of nascent RNA chains (Figure 2A, i). Maximal recruitment would be determined by the size of the transcription unit and would plateau when termination balanced initiation. Previous work established that transcriptional activity during nuclear division cycles of *Drosophila* embryos generally peaks in late interphase and early prophase (Edgar and Schubiger, 1986; Garcia et al., 2013). One would thus predict the gradual growth of the embryonic RNAPII clusters that peaks during late interphase. However, we observed very abrupt emergence of RNAPII clusters shortly after each mitosis (Figure 1A), arguing against this model and suggesting that the accumulation of RNAPII into visible foci can occur independently of transcriptional engagement.

**Figure 2.**
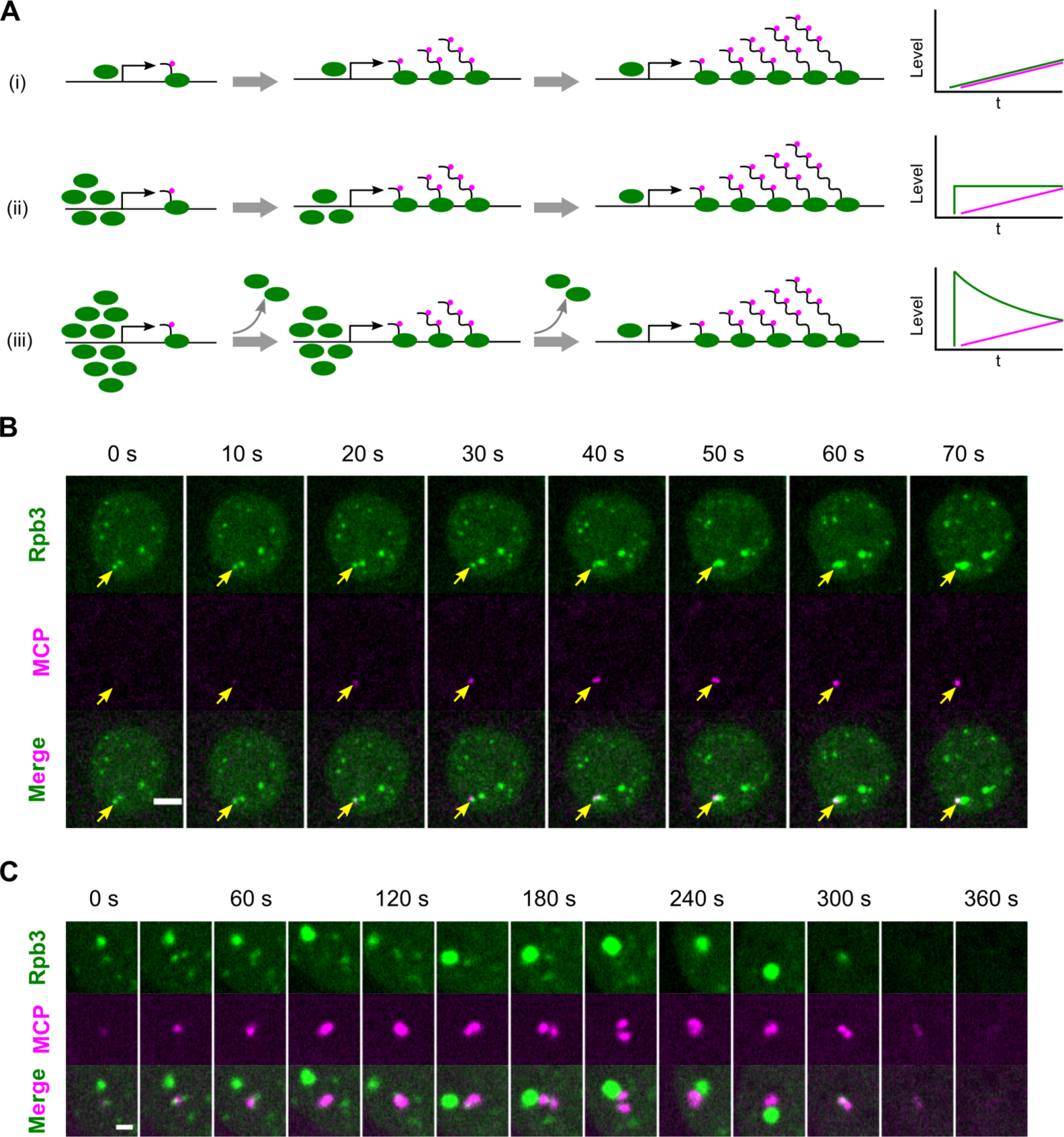
The formation of RNAPII clusters precedes the onset of transcription after mitotic exit. (A) Schematics illustrating possible relationships between the abundances of RNAPII in clusters (green) and of nascent transcript signal (MCP, magenta) during loading of the transcription unit: i) classical recruitment to a promoter; ii) recruitment of an efficiently used pre-transcriptional pool; iii) recruitment of an inefficiently used pre-transcriptional pool. (B) Representative stills from live imaging of EGFP-Rpb3 and MCP-mCherry in an embryo carrying the *hbP2-MS2-lacZ* reporter gene during early nc12. Yellow arrows indicate sites of the same RNAPII cluster colocalizing with MCP spots. The two HLB loci form the brightest Rpd3 foci, and one of these is closely linked and adjacent to the reporter construct: in the late frames it obscures the relevant Rpd3 focus associated with the MCP signal. Maximal projection of 3 z-slices spanning 1 μm is shown. The 0-second timepoint is set as the start of the movie. Scale bar, 2.5 μm. (C) Representative stills from live imaging in the experiments as described in (B) focusing on a single MCP spot going through nc12. Single z-slices with maximal MCP signal are displayed. The 0-second timepoint is set as the start of the movie. A very weak MCP focus at the beginning of the sequence (0 s) aligns with a Rpb3 focus (central), and the large HBL focus of Rpd3 develops next to it. The Rpd3 focus that initially appears coincident with the MCP signal seems to split into two signals bracketing a growing MCP focus (30 s). This is followed by further disruption of the Rpd3 focus and another rearrangement where the MCP signal splits with the Rpd3 sitting centrally (180 s) before it fades. Scale bar, 1 μm. See also Figure S3.

To examine the relationship of RNAPII cluster formation to nascent transcript accumulation more directly, we introduced the MS2/MCP nascent transcript tracking system into embryos expressing EGFP-Rpb3. We focused on a *hbP2-MS2-lacZ* reporter driven by the *hunchback* (*hb*) enhancers and P2 promoter, which is stably expressed in the anterior half of the embryo after nc9 (Garcia et al., 2013). The time of earliest detection of *hbP2-MS2* nascent transcripts (labeled by MCP-mCherry) varied somewhat from nucleus to nucleus in nc11 to nc13 embryos, but always occurred subsequently to the post-mitotic appearance of nuclear RNAPII clusters (Figure S3A and Movie S2). Furthermore, an MCP focus often emerged in apparent association with an RNAPII cluster. When imaged at a frame rate sufficient to track RNAPII clusters, we observed the formation of RNAPII clusters that anticipated appearance of associated MCP foci, in some cases by 10-20 seconds (Figure 2B). The emergence of the nascent transcript signal was often accompanied by an apparent physical disruption of the RNAPII cluster, and the MCP signal seemed to be adjacent rather than comingled with the RNAPII cluster (e.g. Figure 2C, 30-60 seconds). Furthermore, as MCP foci grew larger, the RNAPII clusters shrank but were still juxtaposed to the MCP foci (Figure 2C, 120 seconds). Occasionally, a single small RNAPII cluster was bracketed by two MCP foci, presumably representing transcription on sister chromatids (Figure 2C, 180 seconds). Finally, in late interphase, when MCP foci reached high intensity, associated RNAPII clusters were much reduced or undetectable. We observed a similar outcome using another reporter, *eve2-MS2* (Figure S3B) (Bothma et al., 2014).

These observations show that RNAPII clusters form abruptly prior to significant nascent transcript accumulation, arguing against cluster formation by the direct recruitment of transcriptionally engaged RNAPII (Figure 2A, i). Additionally, the RNAPII cluster does not increase in intensity as the nascent transcript signal increases, further arguing that the observed cluster is not substantially composed of transcribing RNAPII. Nonetheless, the clusters could contribute RNAPII to the transcriptionally active pool as in models outlined in Figure 2A, ii, and iii. Findings described below suggest that enhancer activity and alterations in chromatin state contribute to RNAPII cluster formation, and our imaging shows a cluster closely associated with, but not co-extensive with, a growing focus of nascent transcripts. These observations are consistent with formation of a chromatin-associated pool of RNAPII that promotes and likely contributes to transcription of the adjacent gene. Because the RNAPII cluster fades substantially in advance of dispersal of nascent transcripts, we suggest that only a minority of RNAPII that is recruited to the initial cluster contributes to the pool of transcribing polymerase (Figure 2A, iii).

### Zelda and dCBP promote RNAPII clustering at non-histone genes

A rich literature suggests that the pioneer factor Zelda (Zld) serves an essential role in promoting the minor wave of zygotic transcription in *Drosophila*, yet the precise mechanisms remain unclear. We hypothesized that Zld amplifies transcription through the formation of RNAPII clusters, either through direct interaction or indirectly through local chromatin modifications. To examine whether Zld hubs might directly nucleate RNAPII clustering, we recorded simultaneously the dynamics of sfGFP-Zld and mCherry-Rpb1 (Figure 3). As previously reported, Zld formed dynamic nuclear hubs rapidly after mitosis; during this initial formation period, RNAPII was still uniformly distributed in the nucleus (Figure 3, 0-90 seconds). The Zld foci are more numerous than the later appearing RNAPII clusters. They also appeared to be more short-lived or more mobile, as we can only rarely track individual Zld foci from frame to frame, whereas the majority of RNAPII clusters can be tracked across multiple frames (e.g. Figure 2B). As RNAPII began to form clusters, only a few of them overlapped with Zld hubs (Figure 3, 120-180 seconds), and very few Zld foci had associated Rpb1 signal. These findings indicate that there is not a direct correspondence between Zld foci and the clusters of RNAPII. Nonetheless, it remains possible that a cascade of events transforms some Zld hubs into RNAPII clusters.

**Figure 3.**
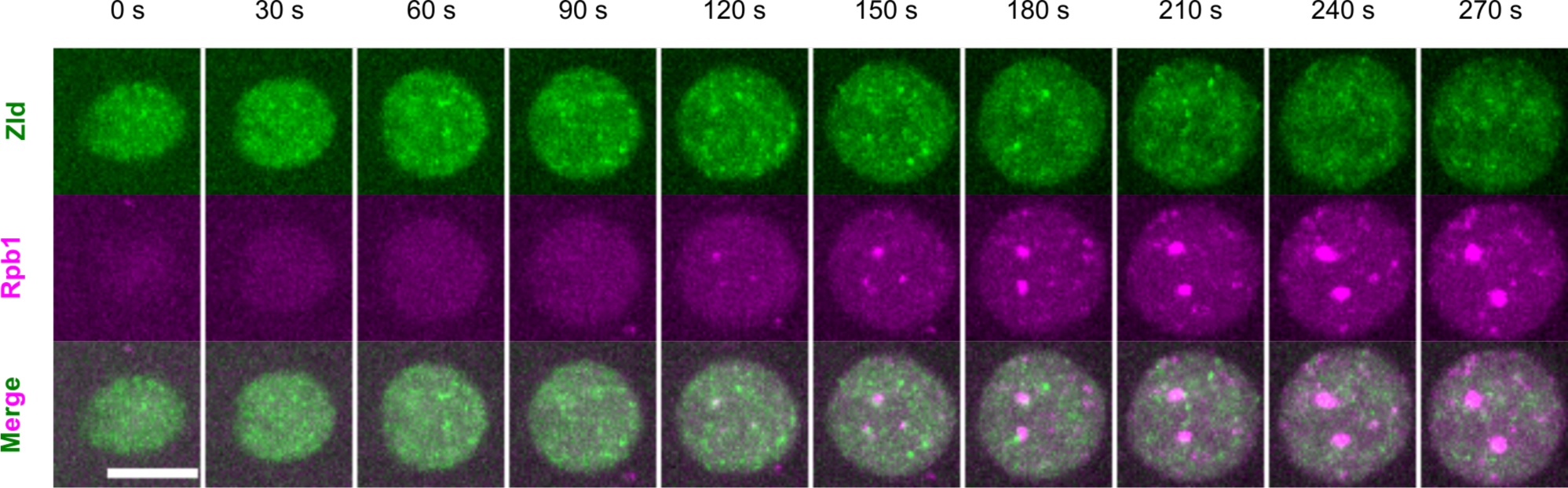
Dynamic clusters of Zelda, a pioneer transcription factor, infrequently and transiently interact with RNAPII clusters. Representative stills from live imaging of sfGFP-Zld and mCherry-Rpb1 in an nc12 embryo. Maximal projection of 7 z-slices spanning 3 μm is shown. The 0-second timepoint refers to the start of this movie. Scale bar, 5 μm.

To test whether the clustering of RNAPII depends on Zld, we employed a JabbaTrap approach to sequester endogenously GFP-tagged Zld in the cytoplasm (Seller et al., 2019) and examined the impact on mCherry-Rpb1 (Figure 4A). The depletion of nuclear Zld using maternally expressed JabbaTrap phenocopied the embryonic lethality the MBT in the *zld* null mutant (Figure S4A). Strikingly, the depletion of nuclear Zld abolished most RNAPII clusters other than the large clusters at HLBs (Figure 4A). This result is consistent with an independent study showing that Zld is required for RNAPII speckles as detected by immunostaining in fixed cells (Huang et al., 2021). The JabbaTrap approach can also be used to abruptly deplete Zld at different nuclear cycles by expressing JabbaTrap from injected mRNA. In past work, we have found that expression of JabbaTrap protein in interphase fails to sequester nuclear GFP-tagged targets until the nuclear membrane breaks down in mitosis. Similarly, we found that interphase accumulated JabbaTrap led to abrupt sequestration of Zld during mitosis. This was accompanied by reduction of RNAPII clustering in the following interphase in nc12 or nc13 (data not shown). This result is consistent with a model that Zld is required to re-establish transcriptional competence after each mitosis during ZGA, but we cannot assess whether Zld is required continuously within each interphase (McDaniel et al., 2019).

**Figure 4.**
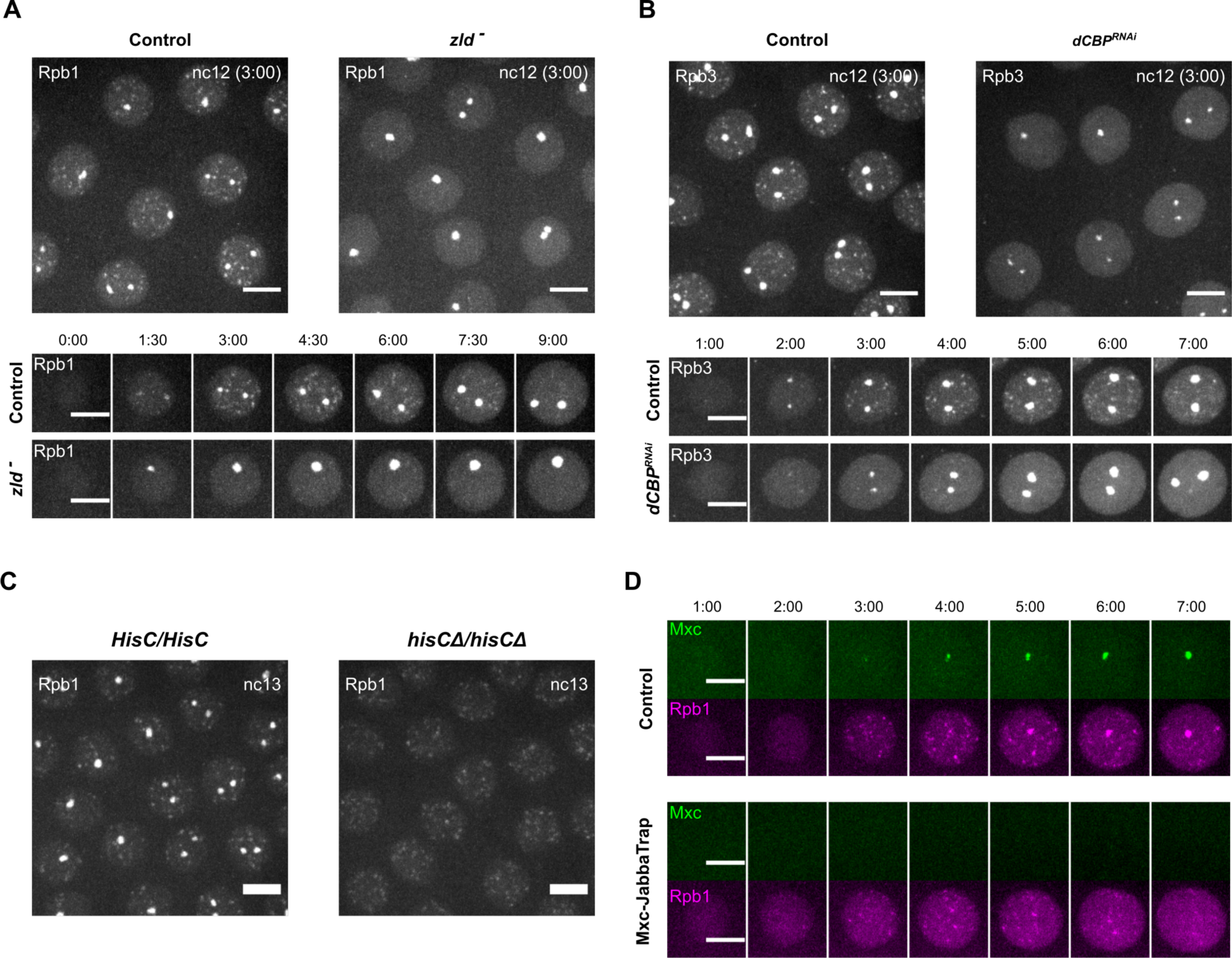
Zelda and the histone acetyltransferase dCBP promote RNAPII clustering at non-histone genes. (A) Representative snapshots from live imaging of mCherry-Rpb1 in the control embryos or the *zld^-^* embryos depleted of nuclear Zelda using a JabbaTrap technique. Both genetic backgrounds maternally expressed the JabbaTrap construct, which is an anti-GFP nanobody fused to the lipid-droplet-binding protein Jabba. In control embryos, the endogenous Zld was untagged and thus unaffected by JabbaTrap. In *zld^-^* embryos, the endogenous Zld was tagged with sfGFP and sequestered in the cytoplasm by JabbaTrap. The depletion of nuclear Zld blocked the formation of RNAPII clusters except the large clusters at HLBs (n ≥ 10 embryos for both control and *zld^-^*). (B) Representative snapshots from live imaging of EGFP-Rpb3 in the control or dCBP/Nej RNAi embryos. Two copies of Mat-tub-Gal4 were used to drive the expression of shRNA against mCherry as control or dCBP/Nejire in the germline. The knockdown of dCBP blocked the formation of small RNAPII clusters (n ≥ 10 embryos for both control and dCBP RNAi). In both (A) and (B), the 0-minute timepoint is set as the first frame with visible nuclear fluorescent signal, which is slightly later than the start of interphase. (C) Snapshots from live imaging of mCherry-Rpb1 in embryos with indicated zygotic genotype. Both embryos are from the *hisCΔ/CyO* females. (D) Representative stills from live imaging of GFP-Mxc and mCherry-Rpb1 in nc12 embryos injected with water (control) or mRNA encoding JabbaTrap to deplete nuclear Mxc. All scale bars in Figure 4 indicate 5 μm. See also Figure S4.

The dependency of RNAPII clustering on Zld despite a failure to observe extensive colocalization suggests an indirect mechanism. Zld binding sites show increased histone acetylation and chromatin accessibility (Harrison et al., 2011; Li et al., 2014; Nien et al., 2011; Schulz et al., 2015; Sun et al., 2015), but the cofactors that work with Zld have not been identified. In other species, the histone acetyltransferase p300/CBP has been found to confer transcriptional competence during ZGA (Chan et al., 2019). The *Drosophila* homolog of CBP is encoded by *Nejire* (*Nej*). The zygotic null *nej* mutant embryos exhibit abnormal zygotic transcription and reduced histone acetylation after the MBT, while germline clones of the *nej* null allele do not produce eggs (Akimaru et al., 1997; Ludlam et al., 2002). We hypothesized that maternally supplied dCBP/Nej mediates histone acetylation at Zld binding sites to promote pre-MBT RNAPII clustering. To knockdown dCBP in early embryos while avoiding sterility, we used a maternal-tubulin-Gal4 driver to express shRNA targeting dCBP during the late stages of oogenesis (Figure S4B)(Park et al., 2019). The dCBP RNAi embryos were inviable and arrested at the MBT (data not shown). Immunostaining for H3K27 acetylation, a modification introduced by CBP that marks active enhancers (Tie et al., 2009), showed a global reduction in dCBP-knockdown embryos (Figure S4C). As with sequestration of Zld in the cytoplasm, RNAi reduction of maternal dCBP blocked most RNAPII clustering in early embryos except the large clusters at HLBs (Figure 4B). We conclude that Zld and dCBP are both required for the formation of RNAPII clusters at non-histone genes. Given that histone acetylation and reduction of nucleosome density are associated with Zld binding (Li et al., 2014; Sun et al., 2015), we suggest that Zld acts upstream of dCBP to initiate local remodeling of chromatin structure and promote RNAPII clustering.

### An Independent pathway controls RNAPII clustering at the histone locus body

The inactivation of Zld or dCBP only affected the formation of small RNAPII clusters but not the large clusters associated with HLBs, showing that a Zld-independent pathway supports RNAPII cluster formation at HLBs. To test whether the small RNAPII clusters require the HLB clusters, we examined embryos carrying homozygous deletions of the *HisC* locus. Removing histone DNA prevented large RNAPII clusters, while the rest of the small clusters still appeared with similar dynamics (Figure 4C). Since RNAPII clustered at the HLB in the absence of nuclear Zld, we hypothesized that a different upstream factor might act at the HLBs. We used JabbaTrap to deplete nuclear Mxc, which is essential for histone gene expression (White et al., 2011). In the absence of nuclear Mxc, RNAPII continued to form small clusters throughout the nucleus without forming the large clusters at HLBs (Figure 4D). Thus, at least two distinct upstream pathways independently regulate RNAPII clustering during *Drosophila* ZGA. One candidate for mediating Zld-independent clustering at the *HisC* locus is CLAMP, a pioneer transcription factor that binds a GAGA repeat in histone gene clusters (Rieder et al., 2017) and is involved in ZGA (Colonnetta et al., 2021; Duan et al., 2021).

### Clustering of transcription machinery and transcript production are prohibited before DNA replication

Since the action of Zld and clustering of RNAPII preceded the onset of transcription, we hypothesized that they might contribute to the post-mitotic temporal delay of transcription as compared to replication. This temporal order could be tightly prescribed if replication served as a necessary precondition for events promoting the onset of transcription. To test this hypothesis, we introduced a block to DNA replication at mitosis and examined the impact on RNAPII clusters and transcriptional activity in the subsequent interphase (Figure 5A).

**Figure 5.**
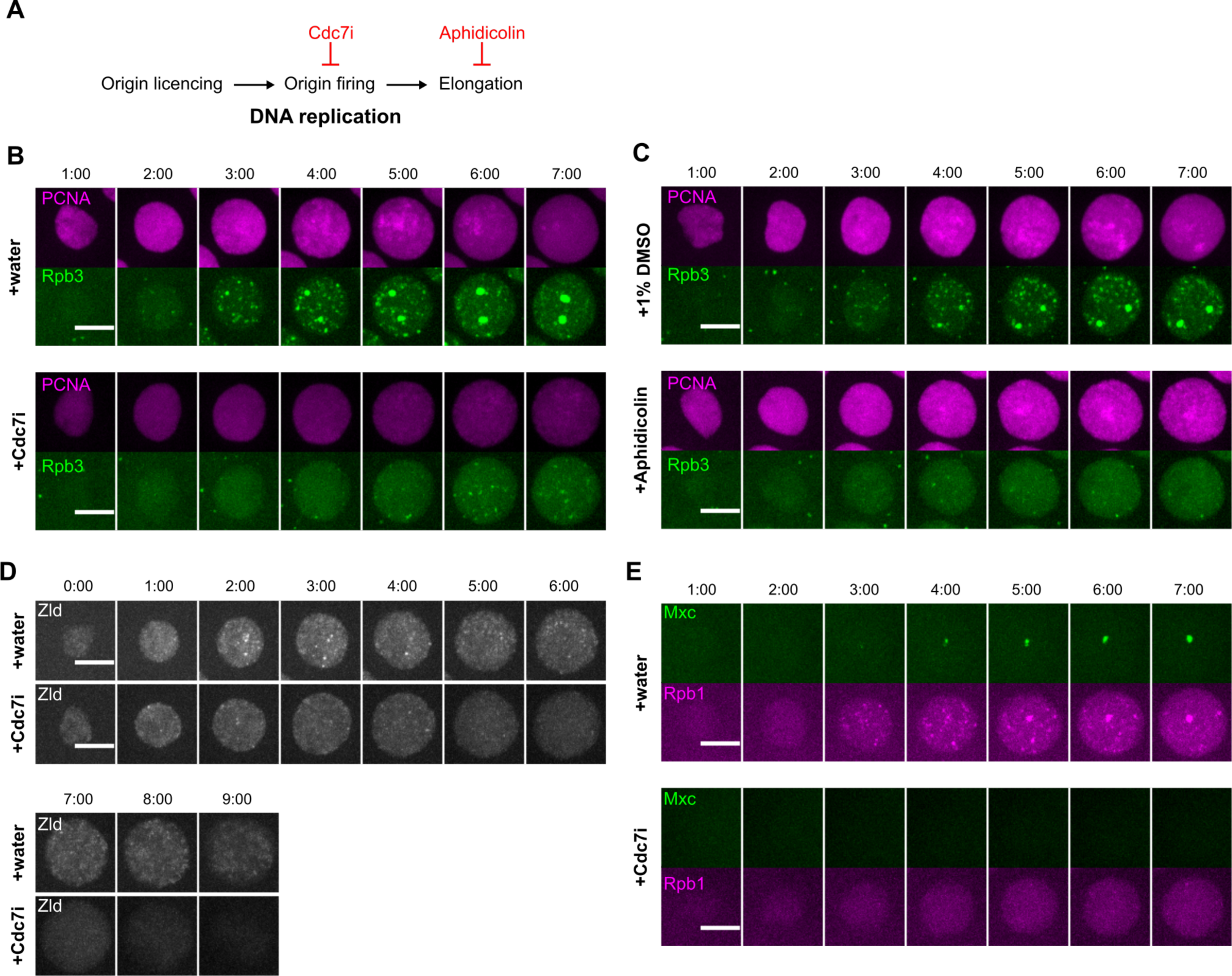
Clustering of RNAPII and its upstream regulators are dependent on DNA replication. (A) Diagram illustrating different steps in DNA replication and corresponding inhibitors. In all experiments in Figure 5 and Figure 6, the injections of inhibitors were performed during mitosis 11, and the following interphase 12 was recorded by confocal live imaging. The injections of control were performed either during interphase or mitosis 11, which yielded similar results. (B) Representative stills from live imaging of mCherry-PCNA, which marks replicating loci, and EGFP-Rpb3 in embryos injected with water or 0.5 mg/ml Cdc7i (n ≥ 3 embryos for both treatments). (C) Representative stills from live imaging of mCherry-PCNA and EGFP-Rpb3 in embryos injected with 1% DMSO as control or 0.1 mg/ml aphidicolin in 1% DMSO (n ≥ 3 embryos for both treatments). In both (B) and (C), the 0-minute timepoint is set as the first frame with visible chromatin-bound PCNA. (D) Representative stills from live imaging of mNeonGreen-Zld in nc12 embryos injected with water or Cdc7i (n = 3 embryos for both treatments). (E) Representative stills from live imaging of GFP-Mxc and mCherry-Rpb1 in nc12 embryos injected with water or Cdc7i (n = 3 embryos for both treatments). The control embryos are the same as those in Figure 4D. All scale bars in Figure 5 indicate 5 μm. See also Figure S5.

We first injected embryos with XL413 (or Cdc7i herein) during mitosis to inhibit Cdc7, an S-phase kinase essential for origin firing (Sheu and Stillman, 2010; Stephenson et al., 2015). Injection during mitosis reduced the signal from a reporter of replication, mCherry-PCNA, and abolished early formation of RNAPII clusters in the next interphase (Figure 5B). Faint RNAPII foci did emerge later in interphase despite Cdc7 inhibition. Nonetheless, the inhibitor clearly delayed and suppressed recruitment of RNAPII. The residual RNAPII cluster formation might simply result from incomplete inhibition of replication or might represent a mechanism different than the early-interphase recruitment of RNAPII. For example, if some transcription initiates in the absence of the RNAPII foci, the eventual accumulation of RNAPII-engaged transcripts on genes ought to produce small foci (Figure 2A, i). We conclude from this result that Cdc7 activity, or some event downstream of its action such as DNA replication, is required for the initial abrupt formation of RNAPII clusters following mitosis.

To test if actual DNA synthesis is required, we injected embryos with aphidicolin, a competitive inhibitor of DNA polymerase alpha. In the injected embryos, the mCherry-PCNA signal and its heterogeneity appeared comparable to control during the beginning of interphase, indicating recruitment of mCherry-PCNA to replication forks (Figure 5C). However, the PCNA signal persisted and was not resolved toward late interphase, consistent with the slowing of the fork progression. Correspondingly, the emergence of RNAPII clusters were largely abolished (Figure 5C). Finally, we injected purified Geminin, a protein inhibitor of Cdt1/Dup and hence origin licensing, and deleted S phase entirely (McCleland et al., 2009). Geminin injection also abolished the RNAPII clusters (Figure S5). We conclude that the formation of RNAPII clusters is tightly coupled to DNA replication in each interphase.

To explore the level of inputs of DNA replication into transcriptional initiation, we next examined the response of upstream transcriptional regulators, including Zld and Mxc, to the inhibition of DNA replication. Surprisingly, we found that inhibiting origin firing by Cdc7i also significantly impaired the formation of Zld and Mxc foci (Figures 5D and 5E), suggesting that DNA replication is a common upstream signal of the two parallel pathways for RNAPII clustering. These results suggest that in these embryonic cycles that lack a G1 phase, DNA replication is required to initiate the observed maturation of nuclear microenvironments and clustering of RNAPII.

These findings suggest the existence of a pre-transcriptional regulatory cascade resulting in RNAPII clustering that depends on replication. If transcription depends on this upstream cascade, it should also depend on replication. To test this dependency, we examined how the inhibition of replication impacted transcription of the *hbP2-MS2* reporter gene. Mitotic injection of Cdc7i delayed and suppressed transcription of the reporter in the subsequent interphase, consistent with a dependency of transcription on prior replication (Figures 6A and 6C). However, several nuclei in these Ccd7i-injected embryos had weak signals of RNAPII clustering and transcriptional activity in late interphase (Figures 6A and 6C). This result could again be explained either by incomplete inhibition of replication by Cdc7i, or by incomplete coupling of transcription to the regulatory cascade. Injection of aphidicolin led to a more stringent block of both RNAPII cluster and transcription throughout interphase (Figures 6B and 6C), arguing that transcription is tightly coupled to the regulatory cascade. Nonetheless, even in the presence of aphidicolin, a few nuclei exhibited a weak transcriptional signal late in interphase. We conclude that the formation of both RNAPII clusters and onset of transcription are coupled to DNA replication, but the evidence of reduced transcription later during interphase suggests that alternate routes of transcriptional initiation might exist.

**Figure 6.**
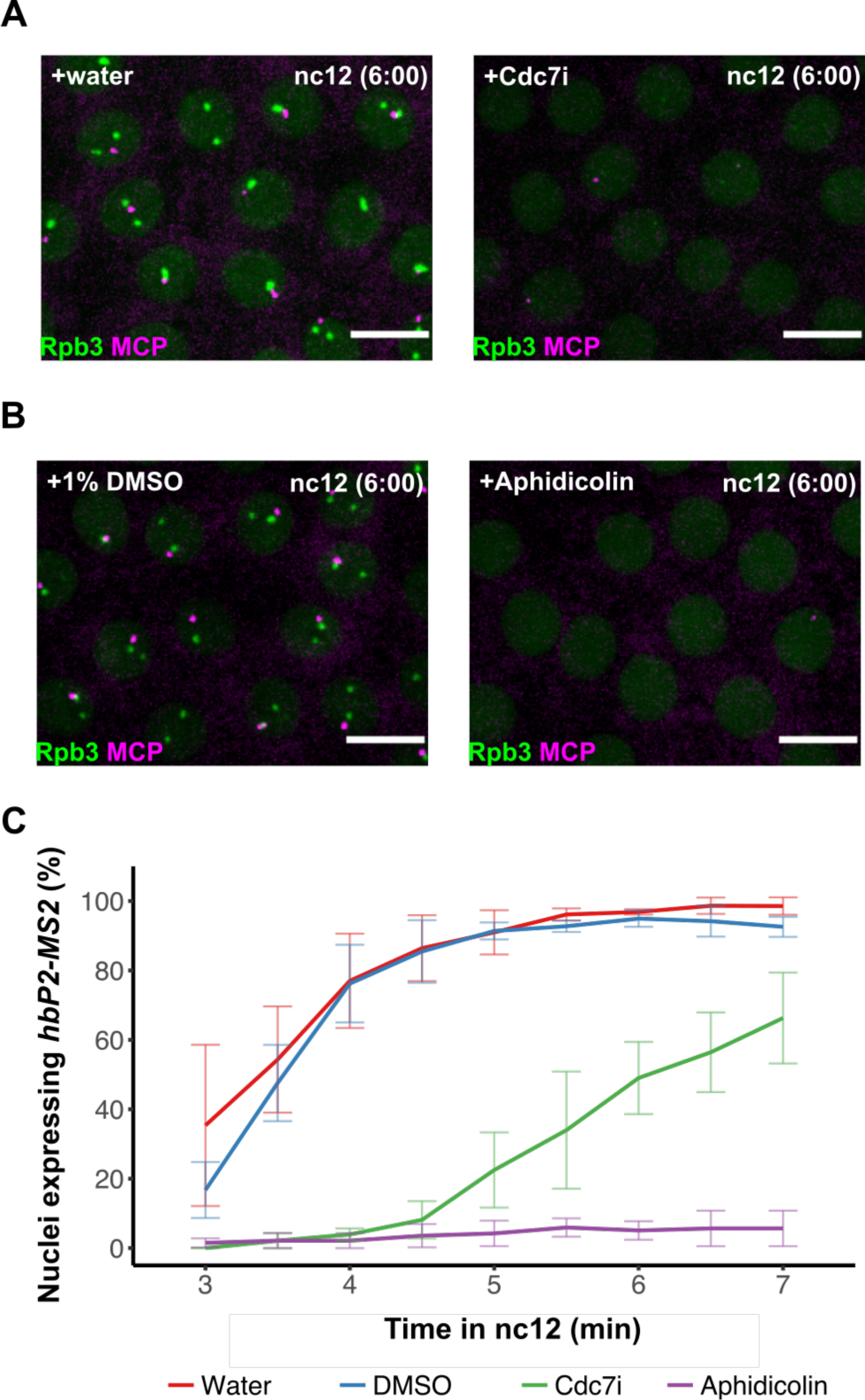
The onset of transcription is dependent on DNA replication. (A and B) Representative stills from live imaging of EGFP-Rpb3 and MCP-mCherry in embryos carrying the *hbP2-MS2-lacZ* reporter (n ≥ 3 embryos for all treatments). The 0-minute timepoint is set as the first frame with visible nuclear EGFP-Rpb3 signal. Scale bars, 5 μm. (C) Percentages of nuclei with MCP foci in embryos carrying *hbP2-MS2* reporter during nc12 after different injections. Note that all detected foci were scored and that the foci detected in the presence of inhibitors were weaker. Error bars represent s.d., n = 3 embryos.

## DISCUSSION

In this study, we have characterized the *in vivo* dynamics of RNAPII nuclear organization during early *Drosophila* embryogenesis, established parallel pathways of RNAPII clustering, and investigated its coordination with the rapid cell cycles. Our data show rapid changes in nuclear microenvironments following mitosis and uncover dependencies indicating that these changes drive the onset of efficient transcription in each successive interphase in early *Drosophila* embryos. Most importantly, we show that these processes are coupled to DNA replication. We propose a model in which the passage of replication forks triggers the sequential clustering of pioneer factors, chromatin modifiers, and RNAPII, thus facilitating the onset of transcription during these extremely short interphases. Moreover, we suggest that the tight coupling of transcriptional initiation to DNA replication minimizes transcription-replication conflicts, thus ensuring both productive transcription and genome integrity (Figure 7).

**Figure 7.**
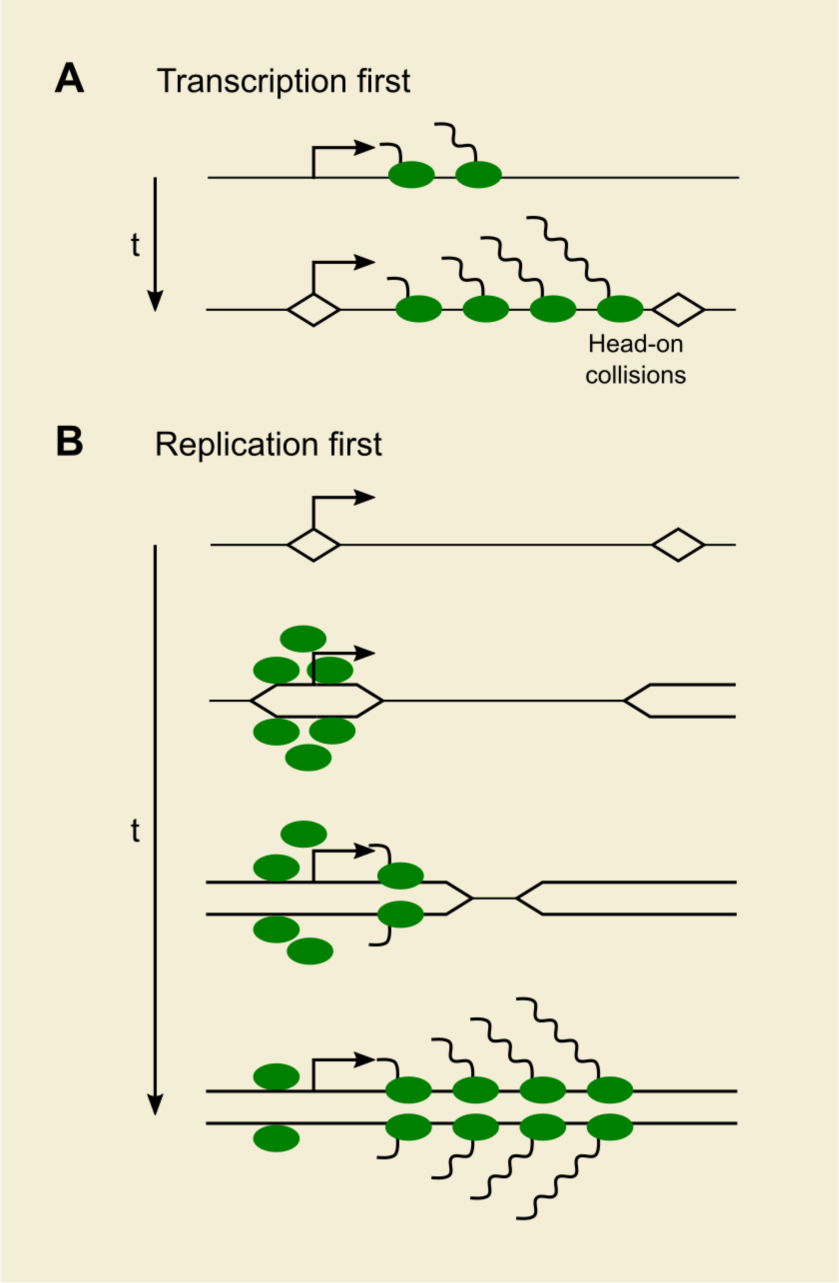
Proposed model for the coordination of transcription and replication in early *Drosophila* embryos. (A) In early *Drosophila* embryos, origins are closely spaced throughout the genome. If transcription initiates before DNA replication, RNA polymerases and replication forks can collide with each other. (B) The embryos resolve this issue by enforcing a temporal order of DNA replication and transcription. The progression of DNA replication facilitates the clustering of RNAPII at promoters and initiates a burst of transcription on sister chromatids behind replication forks. This arrangement avoids conflicts and allows the production of full-length transcripts in short interphases.

In recent years, the clustering or phase separation of transcriptional machinery has emerged as an important mechanism for transcriptional regulation (Cramer, 2019). The formation of liquid-like condensates is thought to enrich a variety of regulators and effectors, including transcription factors, coactivators, Mediator, and RNAPII. Nonetheless, how these nuclear microenvironments are established upon gene activation and regulated during transcription remains an open question. Our work demonstrates the unique advantages of studying transcription in early *Drosophila* embryos and provides several important insights. First, the cell-cycle synchronization of zygotic transcription provides a context in which one can easily follow the time course of events. For example, we show abrupt formation of RNAPII clusters followed by accumulation of nascent transcripts, agreeing with the concepts of “promoter condensates” during transcriptional initiation in mammalian cells (Cramer, 2019). These RNAPII clusters are reduced as transcriptional activity increases as interphase proceeds. This observation might be explained by an RNA-mediated feedback regulation wherein transcriptional condensates are dissolved by high levels of nascent transcripts (Henninger et al., 2021). Second, disruption of gene function can be used to define dependencies in this process. For example, we show a strong dependency of RNAPII clustering on the pioneer factor Zld, even though we could not observe stable association of RNAPII and Zld foci. These data suggest that the initial establishment of transcriptional microenvironments is highly dynamic.

The lack of direct correspondence between Zld foci and RNAPII clustering suggests that the action of Zld requires intermediate steps. In mammalian cells, the BET protein Brd4 binds acetylated histones and is a critical component for phase separation at super-enhancers (Cho et al., 2018). The requirement of the histone acetyltransferase dCBP for RNAPII clustering in *Drosophila* embryos suggests that it might build a target site for recruitment of the Brd4 homolog, encoded by Fsh (Female-sterile homeotic). Furthermore, Zld binding sites exhibit increased histone acetylation (Li et al., 2014; Sun et al., 2015). These observations promote the idea that Zld directs dCBP to acetylate histones near its binding sites, recruiting Fsh to promote RNAPII clustering. Future experiments can test the involvement of Fsh and other candidate factors, while more advanced microscopic techniques should resolve the temporal sequence. In this way, this experimental system should allow a dissection of how the dynamics of nuclear microenvironments might guide gene activation.

The rapid nuclear division cycles pose considerable challenges for early zygotic transcription. Both the short interphase duration and the lack of gap phases significantly limit the window for transcription as compared to a typical somatic cell (Strong et al., 2020). Furthermore, potential conflicts with DNA replication machinery could lead to precocious termination of transcription (García-Muse and Aguilera, 2016). For example, the average distance traversed by replication forks in the early fly embryo is about 4 kb (Blumenthal et al., 1974), a distance similar to the length of many early zygotic genes (Kwasnieski et al., 2019). If transcription were to initiate before replication, widespread head-on collisions could occur (Figure 7A). Linking transcription initiation to prior DNA replication provides an elegant way to tackle these problems. A dependency of RNAPII clustering on prior replication restricts onset of transcription so that RNAPII can follow replication forks without conflicts (Figure 7B). Furthermore, we suggest that replication is followed by processes promoting a transcriptional burst to ensure efficient transcription during the limited temporal window in the early nuclear cycles.

Transcription can certainly occur without DNA replication in G1 or quiescent cells. A wave of post-mitotic transcription in cell lines with gap phases is also well characterized (Hsiung et al., 2016; Palozola et al., 2017). What then makes DNA replication uniquely required for the zygotic transcription in early *Drosophila* embryos? The regulation we describe herein requires two components. First, there must be some constraint that limits transcription on early post-mitotic chromatin. Second, some aspect of replication must reverse or override this constraint. While unknown, we speculate that the relatively naïve genome of early embryos lacking widespread active chromatin marks prevents transcription without DNA replication (Li et al., 2014). One possibility for the reversal of this effect is that increased acetylation levels on early replicated chromatin might act synergistically with Zld and dCBP to open up the chromatin. Secondly, DNA replication might override the limitation by promoting chromatin decompaction after mitosis, as reported for *C. elegans* embryos (Sonneville et al., 2015). Perhaps supporting the latter possibility, inhibiting replication in fly embryos by injecting Geminin results in precocious chromatin condensation in interphase (McCleland et al., 2009). Finally, DNA replication might be involved in establishing chromatin loops and enhancer-promoter contacts, such as those between Zld binding sites (Espinola et al., 2021; Ogiyama et al., 2018).

While the temporal order of DNA replication and transcription provides a basic solution to avoid conflicts, additional coordination might be achieved by placing replication origins close to transcription start sites. The overlap of replication and transcription initiation sites would minimize the lag between the onset of replication at mitotic exit and the later initiation of transcription, thus efficiently using the short interphases. In support of this possibility, replication initiates preferentially from transcriptional start sites of highly transcribed genes in human cells (Chen et al., 2019; Petryk et al., 2016). In both *Drosophila* cells and *C. elegans* embryos, active chromatin marks and highly transcribed genes correlate with replication initiation sites (Comoglio et al., 2015; Pourkarimi et al., 2016; Rodríguez-Martínez et al., 2017). In this regard, it would be interesting to test whether Zelda binding sites correlate with replication initiation sites. Notably, zygotic genome activation has been proposed to generate transcription-replication conflicts, activate the DNA replication checkpoint, and lead to S-phase lengthening in early fly embryos (Blythe and Wieschaus, 2015). Nonetheless, the direct evidence that stalled RNAPII collides with replication forks and slows down the cell cycle is still lacking. For example, Zld-dependent events upstream of RNAPII recruitment may be responsible for changing the global replication-timing program around the MBT. Further work is needed to fully understand the interplay between zygotic genome activation and S-phase reprogramming in early embryogenesis.

Because the early embryonic wave of gene expression is uniquely exposed to the acute threat of collisions with replication forks, we cannot exclude the possibility that the temporal ordering of replication and transcription evolved only to avert this special threat. Alternatively, the *Drosophila* embryos may have adopted and optimized a general mechanism to limit transcription to a short but favorable cell cycle window. The transition from compacted anaphase chromosomes to onset of replication and to initiation of transcription is remarkably fast, taking about 1 and 3 minutes respectively. Other systems have not been analyzed at these time scales, so we know little about the speed and ordering of events during this dramatic transformation. The specialization introduced in the fly embryo might be to use the incredibly rapid onset of replication to accelerate and coordinate onset of transcription. The slower cell cycles in other systems follow a more relaxed time course. For example, a post-mitotic telophase spike in transcription has been detected in mammalian cells, but this occurs about an hour after release from a metaphase block (Hsiung et al., 2016). While there may be no need for replication input in such cell cycles, upon S-phase entry, mechanisms are still needed to carefully coordinate transcription and replication, such as for the replication-dependent histone genes. While recent work indicates that the known limitations for transcription of the histone repeats are satisfied at the restriction point in the mammalian cell cycle, full activation of transcription still awaits the onset of replication (Armstrong and Spencer, 2021). Since histone genes need to be heavily transcribed early in S phase to meet demand, avoidance of collisions is likely to be important, and perhaps mammalian histone gene expression will share the pathway used to activate zygotic transcription in *Drosophila*.

## ACKNOWLEDGEMENTS

We thank the Bloomington Drosophila Stock Center for providing fly stocks. We thank Michael Stadler and Michael Eisen for kindly sharing the mNeonGreen-Zld fly stock and for communicating their early results on RNAPII clustering. We thank members of the O’Farrell lab, Pei-I Tsai and Ekaterina Korotkevich, for critical reading and helpful discussion. This work is supported by National Institutes of Health grants R01GM058921 (to R.J.D) and R35GM136324 (to P.H.O).

## AUTHOR CONTRIBUTIONS

**C.-Y.C.** Conceptualization, Methodology, Formal analysis, Investigation, Writing – Original Draft, Visualization. **J.P.K.**: Methodology, Validation, Resources, Writing – Review & Editing. **R.J.D**.: Resources, Writing – Review & Editing, Supervision, Funding acquisition. **P.O’F**: Conceptualization, Writing – Original Draft, Supervision, Funding acquisition.

---

**Figure S1, Related to Figure 1.**
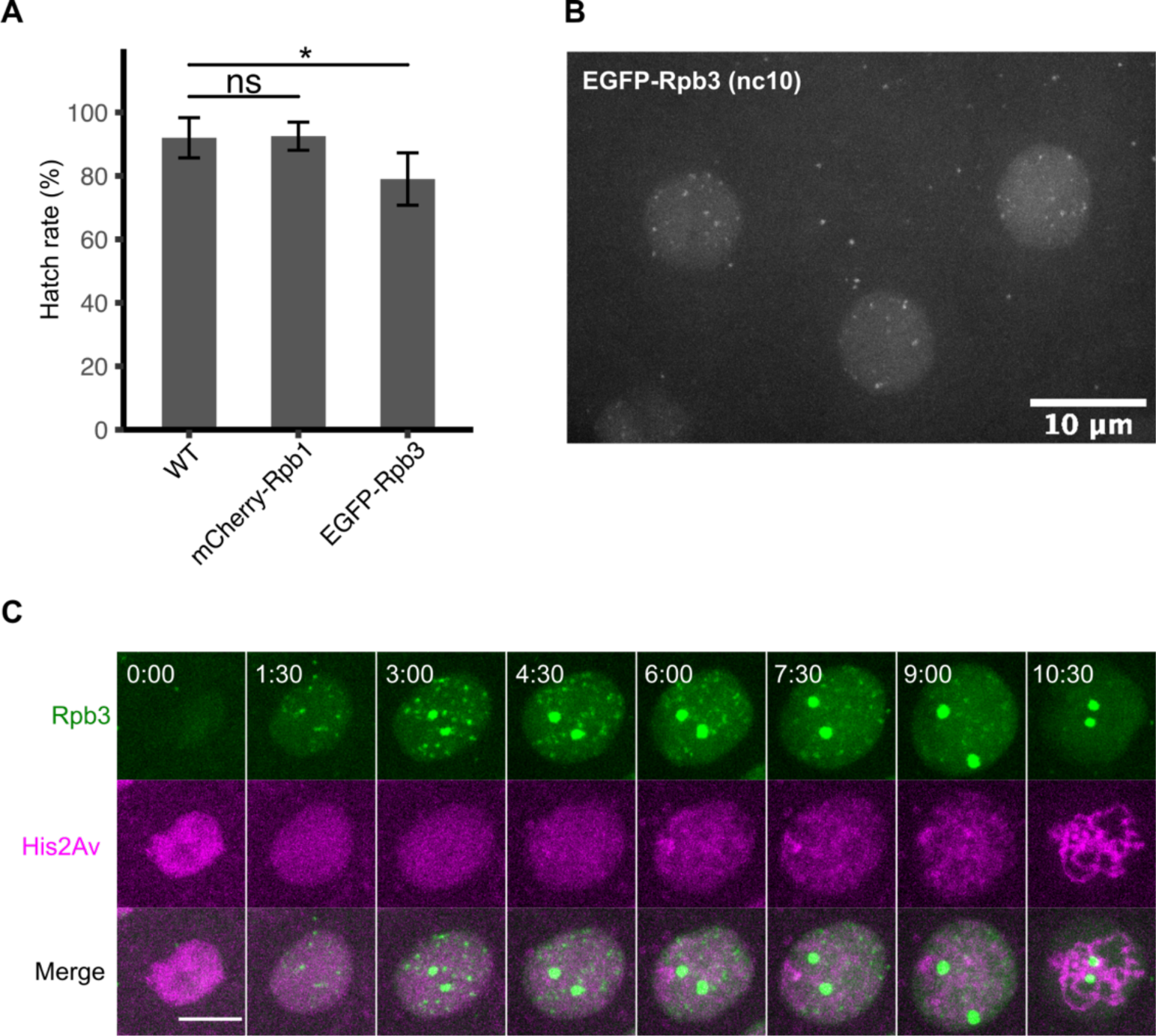
Endogenously tagged RNA polymerase II (RNAPII) subunits form dispersed nuclear foci in syncytial-blastoderm stage embryos of *Drosophila*. (A) Embryo hatch rates in wild type or fly lines carrying endogenously tagged RNA polymerase II subunits. The genotypes indicate both that of mothers and embryos. Data are shown as mean ± s.d. (n = 4 independent experiments). *p<0.05 by one-tailed t-test. (B) A snapshot of an embryo expressing EGFP-Rpb3 at nc10, when the nuclei first migrated to the surface of the embryo. (C) Representative stills from live imaging of EGFP-Rpb3 and His2Av-RFP in embryos during nc12. The timepoint 0 is set as the first frame when nuclei marked by His2Av changed into oval shape after mitosis. Scale bar, 5 μm

**Figure S2, Related to Figure 1.**
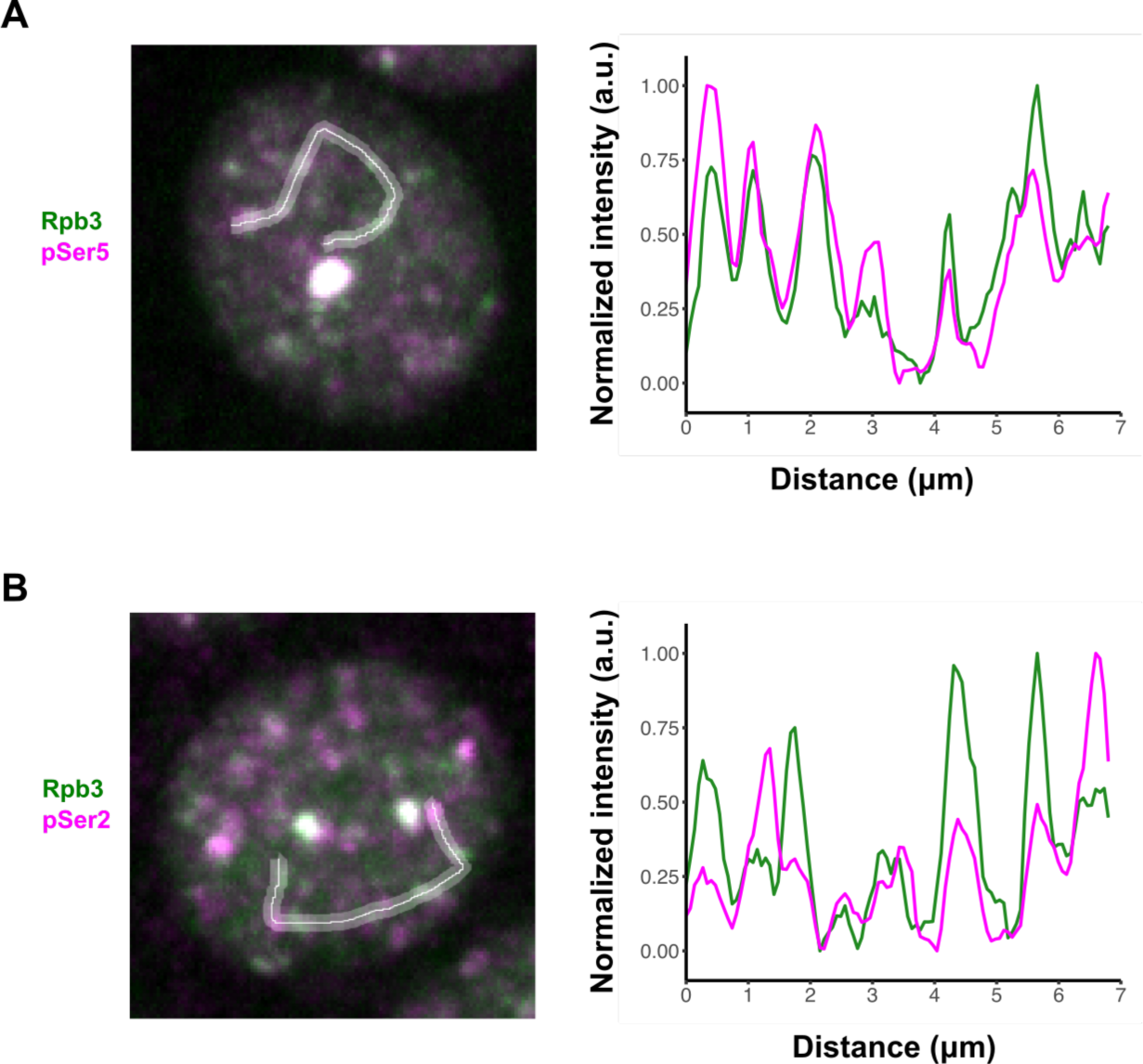
Relationships between Rpb3 foci and immunostaining foci for pSer5 or pSer2. Intensity profiles (right) were measured from freehand lines as shown on left. Images are the same as shown in Figures 1D and 1E.

**Figure S3, Related to Figure 2.**
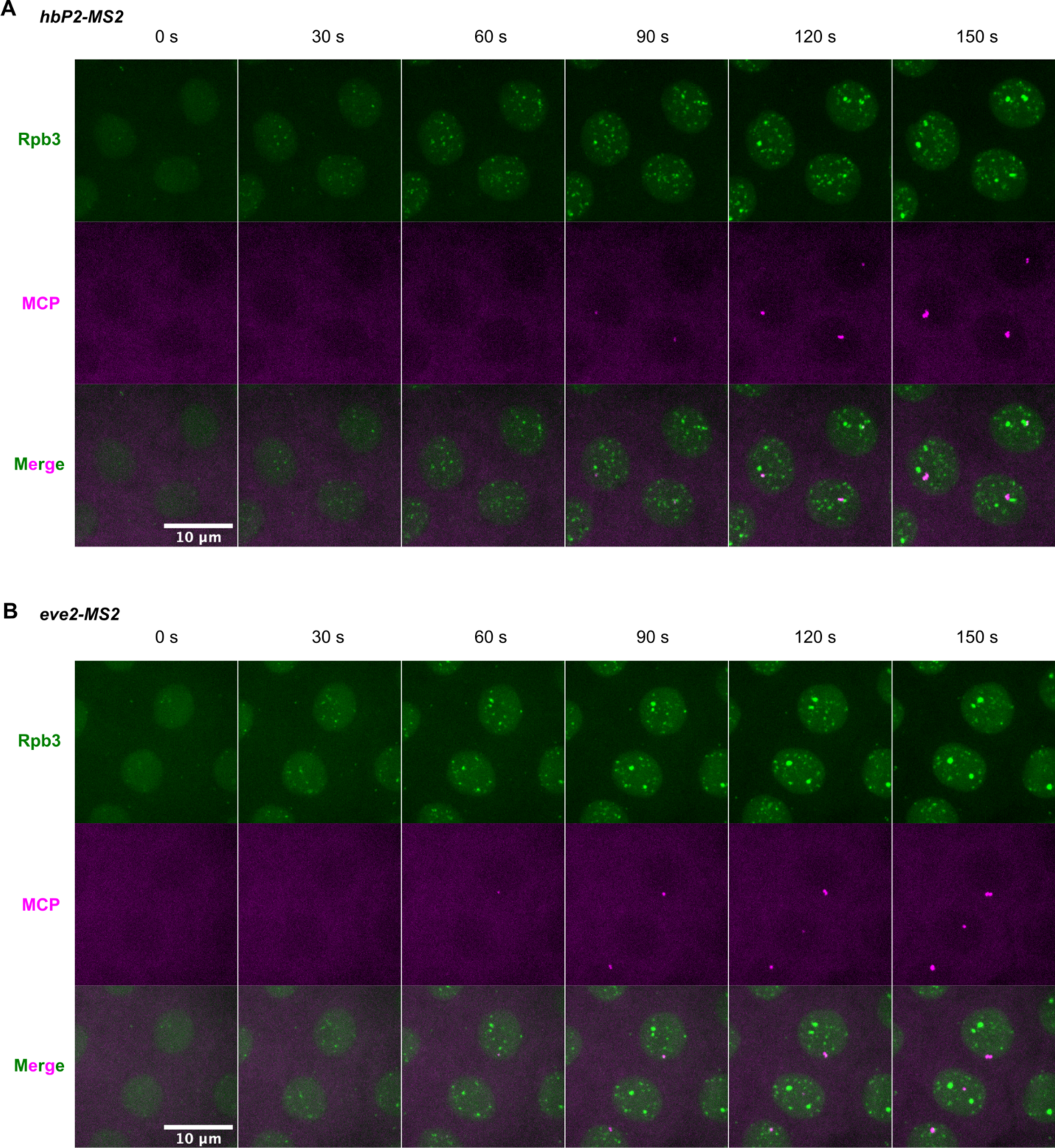
Live imaging of RNAPII clusters and MS2 transcriptional reporters. Representative stills from live imaging of EGFP-Rpb3 and MCP-mCherry in embryos carrying hbP2-MS2 (A) or eve2-MS2 (B) reporters during nc12.

**Figure S4, Related to Figure 4.**
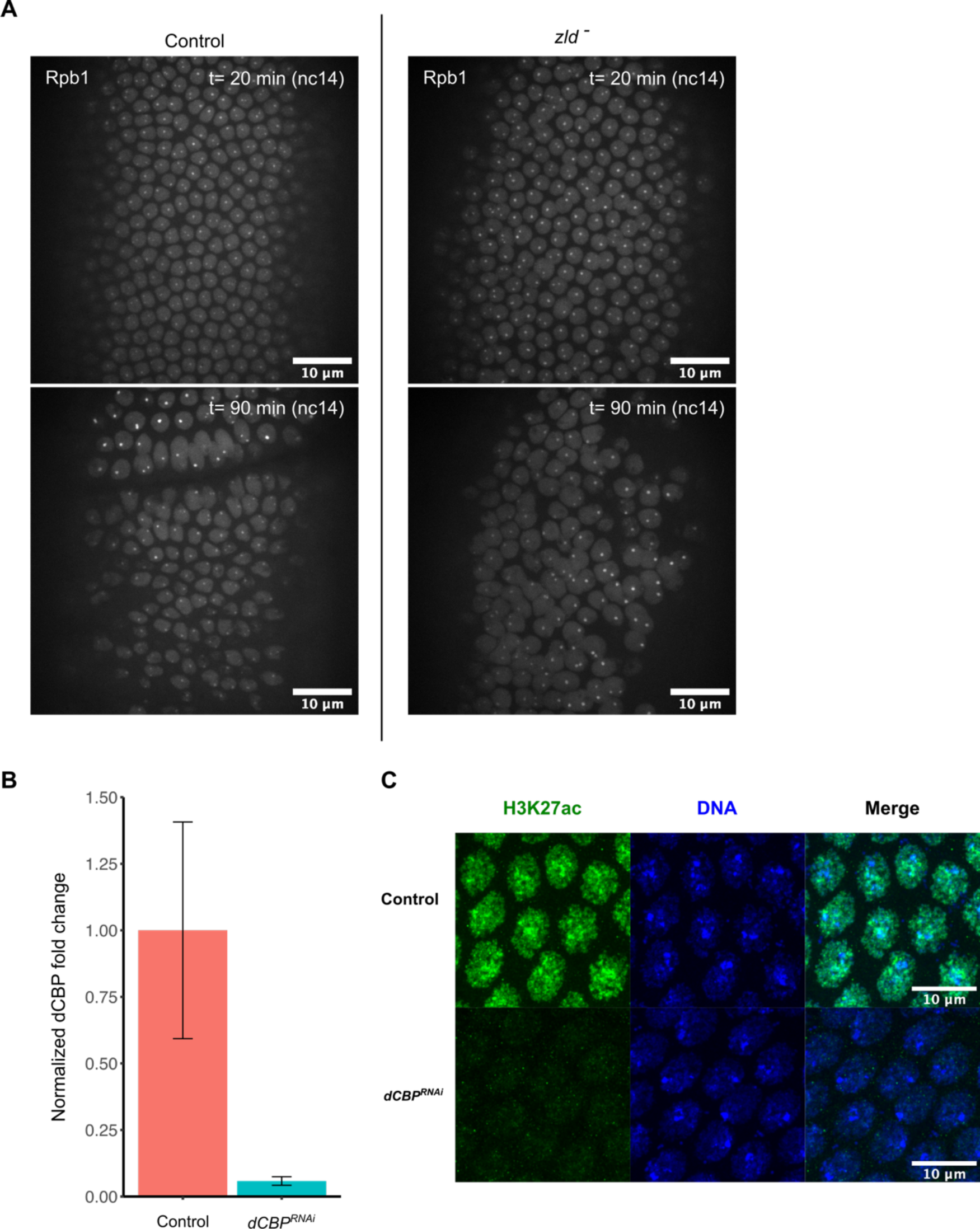
Additional information of embryos with Zld or dCBP inhibitions. (A) Representative still from live imaging of mCherry-Rpb1 in control or JabbaTrap (*zld^-^)* embryos during nc14 at the MBT. The cephalic furrow can be seen in control embryos at 90 minutes, whereas the *zld^-^*embryos fails at both cellularization and gastrulation. (B) Fold change of *dCBP* transcript levels normalized by the reference gene *RpL32* in 0-1.5-hour old embryos. Error bars represent s.d., n = 3 biological replicates. (C) Immunostaining of control and dCBP RNAi nc13 embryos against H3K27 acetylation and DNA. Images are displayed with the same normalization of intensity.

**Figure S5, Related to Figure 5.**
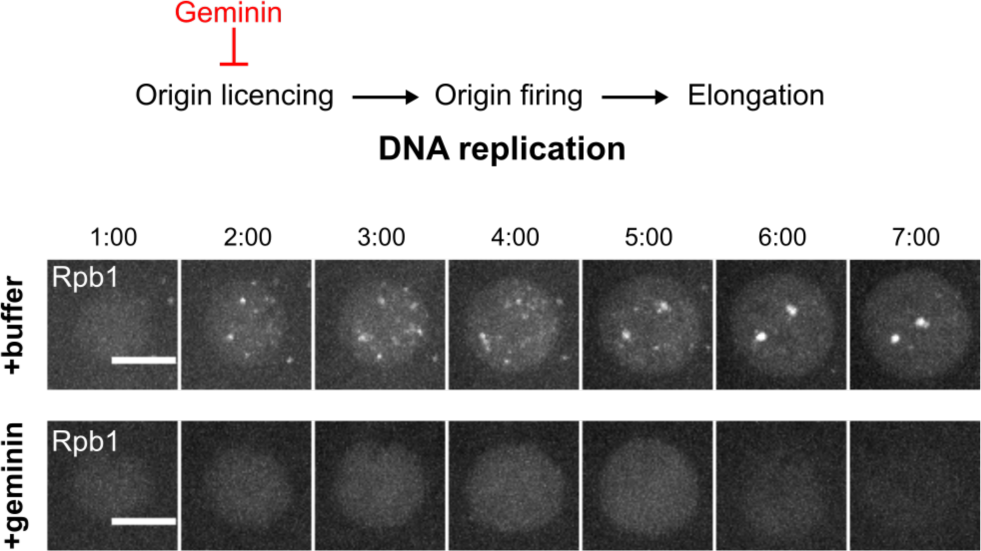
Inhibiting DNA replication impairs RNAPII clustering. Representative stills from live imaging of mCherry-Rpb1 in nc12 embryos injected with buffer or geminin. Scale bar, 5 μm.

## STAR Methods

### RESOURCE AVAILABILITY

#### Lead contact

Further information and requests for resources and reagents should be directed to the lead contact, Patrick H. O’Farrell.

#### Material availability

Fly stocks generated in this study are available from the lead contact upon request.

#### Data availability

The raw imaging data and custom scripts are available from the lead contact upon request.

### EXPERIMENTAL MODEL AND SUBJECT DETAILS

#### Fly stocks

All experiments were performed on *Drosophila melanogaster*. Fly stocks were maintained on standard cornmeal-yeast medium at 25°C. Flies were transferred to egg-laying cages 2-3 days before experiments, and embryos were collected on grape juice agar plates. Fly lines used in this study are listed in the Key Resources Table.

### METHOD DETAILS

#### Molecular cloning and CRISPR-Cas9 genome editing

Oligos for sgRNA targeting 5’ ends of Rpb1 and Rpb3 were ordered from IDT, annealed and cloned into pU6-BbsI-chiRNA using standard protocol. To make donor plasmids, about 1 kb of homology arms upstream and downstream of start codons were amplified from genomic DNA of the *vas-Cas9* flies. EGFP and mCherry DNA fragments with 5xGGS linker were amplified from plasmids previously made in the lab. The pDsRed-attP was cut with XhoI and HindIII to be used as vector backbone. All DNA fragments were then purified by gel purification and assembled by Gibson assembly. The sgRNA and donor plasmids were sent to Rainbow Transgenic Flies (Camarillo, CA) for microinjection. After injection, surviving adults were crossed to appropriate balancer stocks and screened by PCR for successful knock-in. Transformants were backcrossed with wild type at least three times before performing experiments.

#### Hatch rate assay

Embryos were collected for 2-4 hours in egg-laying cages, washed with water, and transferred to grape juice agar plates. 50 embryos in each assay were aligned in groups of 10. Percentages of embryos hatched were then scored after 36 hours of incubation at 25°C.

#### Spinning disk confocal microscopy

Imaging of live or fixed embryos was performed on an Olympus IX70 microscope equipped with PerkinElmer Ultraview Vox confocal system. All experiments were performed at room temperature with 63x or 100x oil objective, and binning was set to 1x1 or 2x2 depending on the experiments. Data were acquired using Volocity 6 software (Quorum Technologies). Fluorophores were excited with 405, 488, and 561 nm laser lines and imaged with proper emission filters. Focal planes each 0.5-0.75 μm apart were recorded at each timepoint, spanning 3-10.5 μm across the nuclei at the surface of the embryos. For experiments in Figures 1B, 1C, 2C, and 3, images were acquired in the order of channels first and then z-planes. In the rest of experiments, images were acquired in the order of z-planes first and then channels. Images in the same set of experiments were acquired using the same configuration, and laser power was calibrated using a laser power meter (Thorlabs) before each imaging session.

#### Embryo fixation and immunostaining

##### Formaldehyde fixation

Embryos were collected in egg-laying cages, washed and transferred into a basket, dechorionated with 25% bleach for 2 minutes, washed three times with water, and transferred into a 1.5-ml tube with 500 μl heptane. Samples were then added with 500 μl of fresh 4% formaldehyde in PBS and then vigorously shaken for 20 minutes at room temperature. After removing the lower aqueous layer carefully, 500 ul methanol was added, and the mixture was shaken for 1-2 minutes. Devitellinated embryos that sank to the bottom were kept and washed with methanol for three times. Fixed embryos were stored frozen at -20°C in methanol until use.

##### Immunostaining

Fixed embryos were rehydrated with 500 ul PBST (0.3% Tween-20) for 5 minutes at room temperature four times. Embryos were then blocked in PBST with 3% donkey serum for 30 minutes. Primary antibodies were added and incubated at 4°C overnight. Embryos were then washed with PBST for 15 minutes three times, incubated with secondary antibodies at room temperature for 1 hour, washed with PBST for 15 minutes three times, and mounted on a glass slide in Fluoromount. All primary and secondary antibodies were used at 1:500. Hoechst 33258 was used at 1:2000 during the second wash after secondary antibodies. Antibodies used are listed in the Key Resources Table.

#### Embryo RNA extraction and RT-qPCR

For each biological replicate, about 100 dechorionated embryos were homogenized in TRI Reagent, and the total RNA was extracted using Direct-zol RNA microprep kit (Zymo Research, #R2060). cDNA was synthesized using Promega GoScript Oligo(dT) (#A2790) following manufacturer’s protocol. qPCR mixture was prepared using Bio-Rad SsoAdvanced Universal SYBR Green Supermix (#172-5270) and run on Bio-Rad CFX Connect system. *RpL32* was used as a reference gene, and dCBP level was quantified by the ΔΔCt method.

#### Embryo mounting for live imaging

Embryos were collected in egg-laying cages, washed and transferred into a basket, dechorionated with 25% bleach for 2 minutes, washed three times with water, and transferred onto grape juice agar plates. Embryos were aligned and transferred to coverslips with glue derived from double-sided tape using heptane. The embryos were then covered with halocarbon oil (1:1 mixture of halocarbon oil 27 and 700) for experiments.

#### Microinjection

After aligned and glued to a coverslip, embryos were desiccated in a desiccation chamber for 7-9 minutes before being covered in halocarbon oil for microinjection. XL413 was dissolved in water at 5 mg/ml and stored at -20°C as stock solution. Aphidicolin was dissolved in DMSO at 10 mg/ml as stock solution. Injections were performed at the following concentrations: 0.5 mg/ml XL413/Cdc7i, 0.1 mg/ml aphidicolin (in 1% DMSO), and 20 mg/ml ECFP-geminin (in 40 mM HEPES pH 7.4, 150 mM KCl buffer).

#### Fly crosses for MS2/MCP imaging

Virgins of EGFP-Rpb3 (II); nos-MCP-mCherry (III) were crossed with males of *hbP2-MS2-lacZ* for 2-3 days. Flies were then transferred together into a cage, and embryos laid were collected for imaging. All crosses and embryo collection were performed at 25°C.

#### Fly crosses for JabbaTrap experiments

The cages for JabbaTrap were set up in two crosses. First, virgins with UASp-JabbaTrap were crossed with Mat-tub-Gal4 males with the same fluorescent reporters. The resulting progenies, either mCherry-Rpb1 (X);; Mat-tub-Gal4/UASp-JabbaTrap (III) or mCherry-Rpb1, sfGFP-Zld (X);; Mat-tub-Gal4/UASp-JabbaTrap (III), were crossed with their siblings and transferred to cages for experiments. To increase the Gal4/UASp expression, F1 were grown at 27°C since larval stage, and cages were also kept at 27°C for embryo collection.

#### Fly crosses for germline RNAi experiments

The cages for RNAi were set up in two crosses. First, virgins of EGFP-Rpb3 (II); UASp-shRNA (III), where the shRNA targeted either mCherry or dCBP/Nej, were crossed to males of EGFP-Rpb3, Mat-tub-Gal4 (II); Mat-tub-Gal4 (III). The resulting progenies were then crossed with their siblings and transferred to cages for experiments. To increase the Gal4/UASp expression, F1 were grown at 27°C since larval stage, and cages were also kept at 27°C for embryo collection.

#### Image processing and presentation

Data obtained in Volocity were exported as image stacks and processed in FIJI/ImageJ and Python. Maximal projections are shown in figures unless otherwise noted.

### QUANTIFICATION AND STATISTICAL ANALYSIS

#### Quantification of the percentage of nuclei transcribing hbP2-MS2 reporter

Analyses were performed on maximal projections from live imaging of embryos expressing EGFP-Rpb3 and MCP-mCherry with a *hbP2-MS2-lacZ* reporter. To identify MCP spots, images of MCP-mCherry were first background-subtracted with a rolling ball radius of 25 pixels in FIJI, and then the Trainable Weka Segmentation tool was used to detect MCP foci. The “FastRandomForest” classifier was trained on selected frames from one movie of an embryo injected with water and then run on all data. Segmentation results were exported and used for downstream analyses by custom Python scripts. EGFP-Rpb3 images were used to identify and count the number of nuclei. The timepoint 0’ was determined as the first frame with visible EGFP-Rpb3 nuclear signal. Because of the weak EGFP-Rpb3 intensity at the beginning of interphase, we performed analyses on data beginning at 3 minutes. Briefly, EGFP-Rpb3 images were blurred by Gaussian filter and then converted to binary masks using otsu thresholding. After filtering out small nuclei at the edge of images, the number of total nuclei and those contained MCP spots were counted, and percentages of nuclei transcribing the reporter were calculated.

---

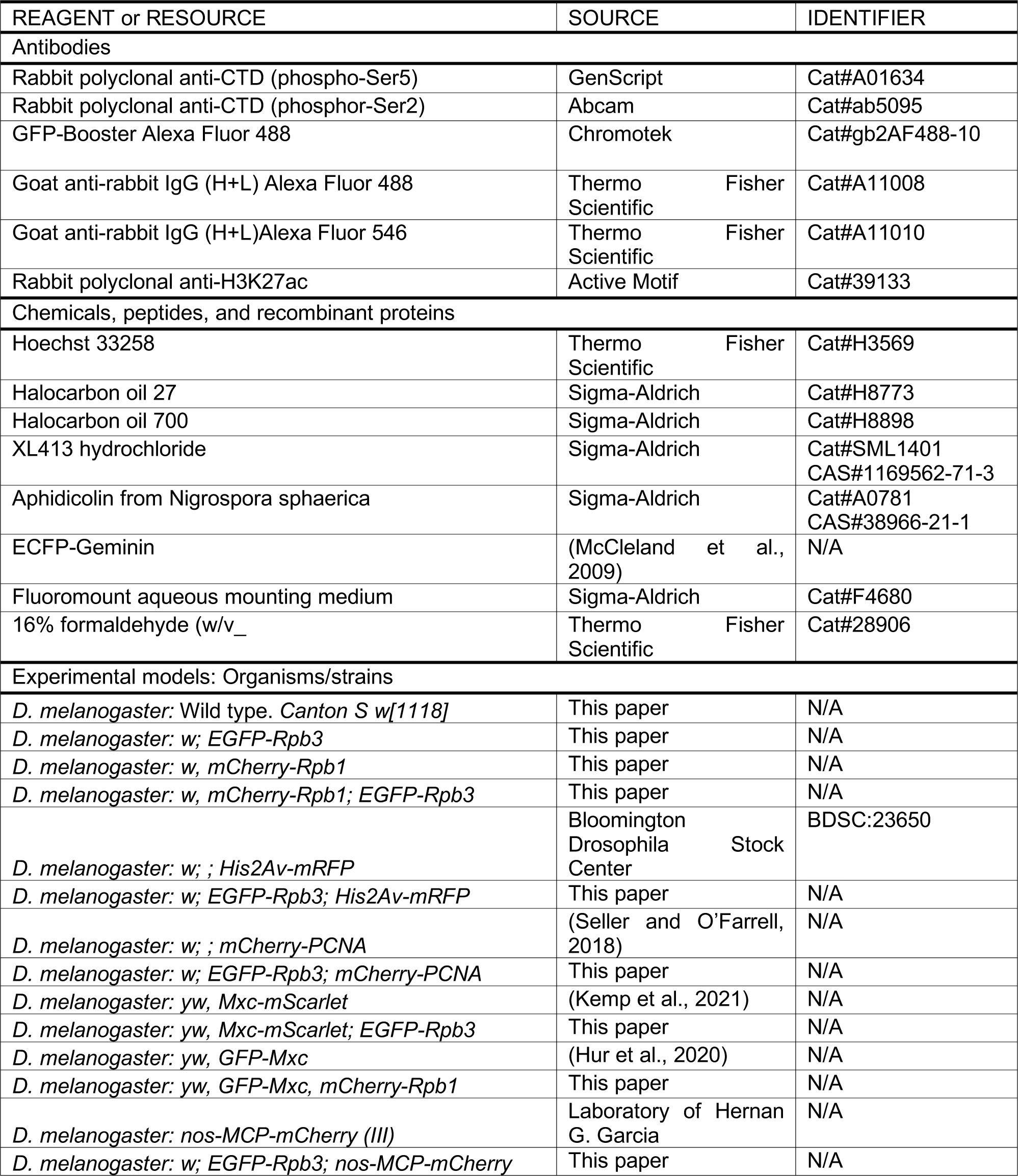
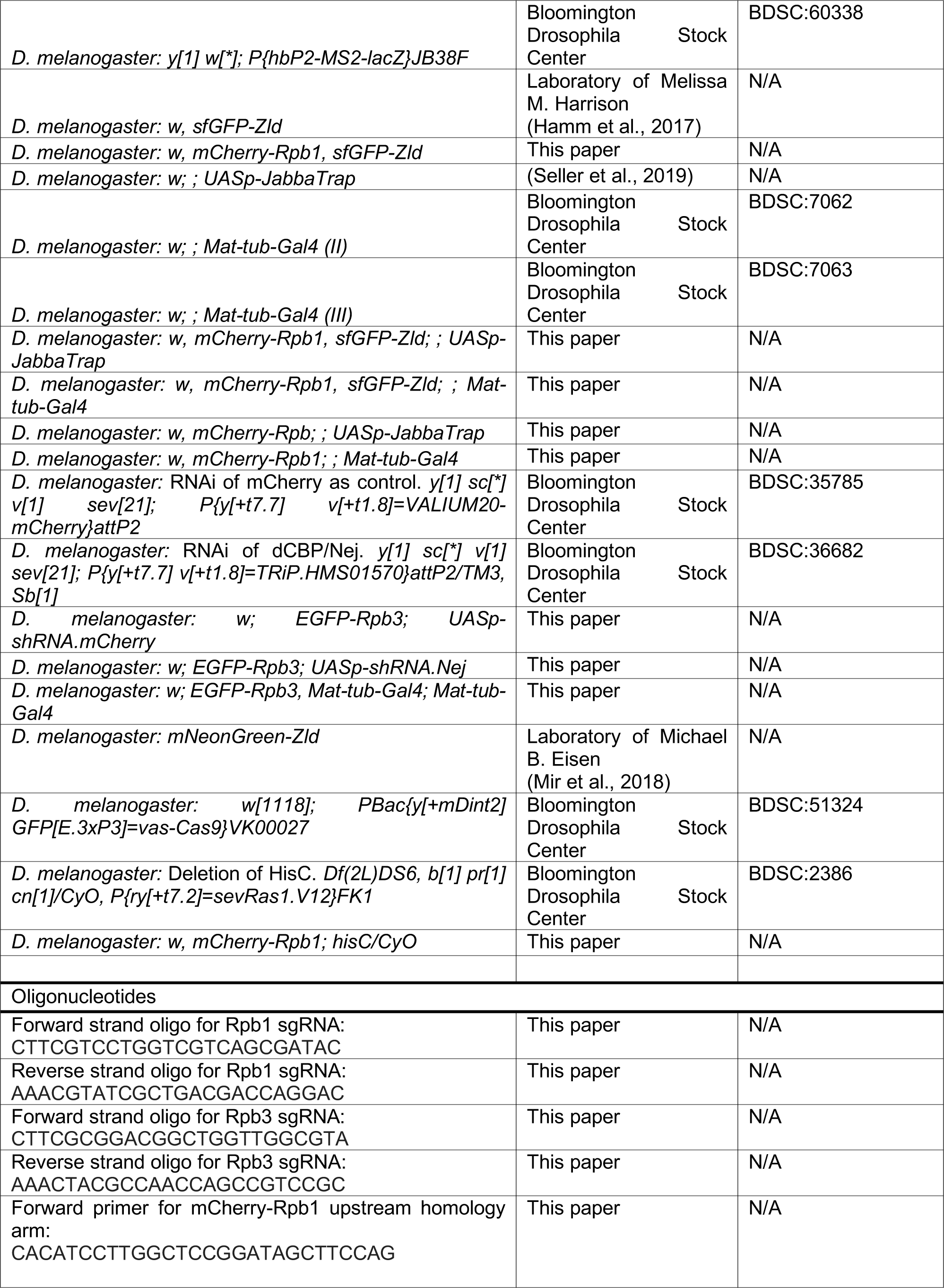
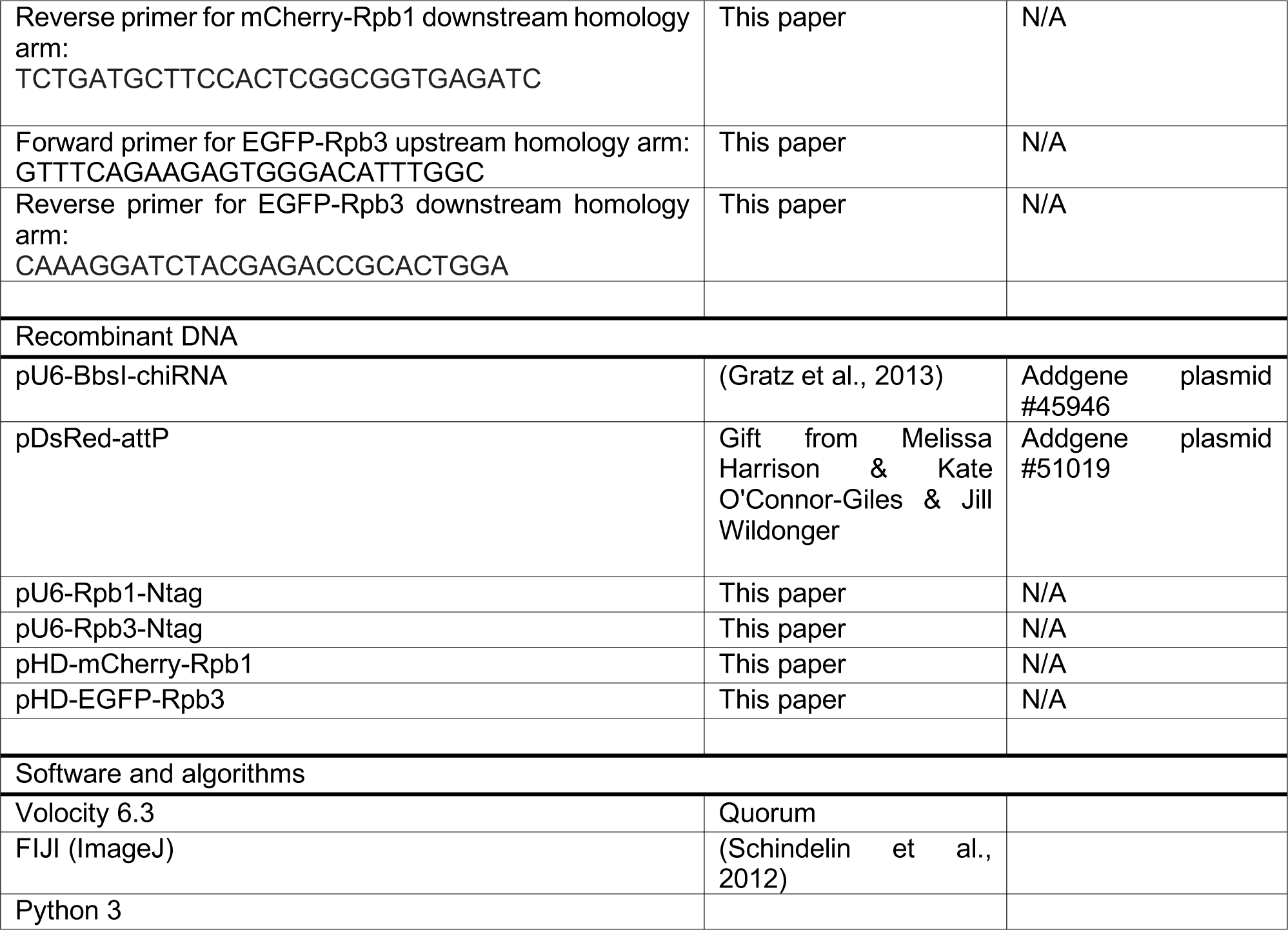
KEY RESOURCES TABLE.

## REFERENCES

1. Akimaru, H., Hou, D.X., and Ishii, S. (1997). Drosophila CBP is required for dorsal-dependent twist gene expression. Nat. Genet. 17, 211–214.

2. Armstrong, C., and Spencer, S.L. (2021). Replication-dependent histone biosynthesis is coupled to cell-cycle commitment. PNAS 118, 2100178118.

3. Bedinger, P., Hochstrasser, M., Victor Jongeneel, C., and Alberts, B.M. (1983). Properties of the T4 bacteriophage DNA replication apparatus: The T4 dda DNA helicase is required to pass a bound RNA polymerase molecule. Cell 34, 115–123.

4. Blumenthal, A.B., Kriegstein, H.J., and Hogness, D.S. (1974). The units of DNA replication in Drosophila melanogaster chromosomes. Cold Spring Harb. Symp. Quant. Biol. 38, 205–223.

5. Blythe, S.A., and Wieschaus, E.F. (2015). Zygotic genome activation triggers the DNA replication checkpoint at the midblastula transition. Cell 160, 1169–1181.

6. Blythe, S.A., and Wieschaus, E.F. (2016). Establishment and maintenance of heritable chromatin structure during early Drosophila embryogenesis. Elife 5, e20148.

7. Boehning, M., Dugast-Darzacq, C., Rankovic, M., Hansen, A.S., Yu, T., Marie-Nelly, H., McSwiggen, D.T., Kokic, G., Dailey, G.M., Cramer, P., et al. (2018). RNA polymerase II clustering through carboxy-terminal domain phase separation. Nat. Struct. Mol. Biol. 25, 833–840.

8. Boija, A., Klein, I.A., Sabari, B.R., Dall’Agnese, A., Coffey, E.L., Zamudio, A. V., Li, C.H., Shrinivas, K., Manteiga, J.C., Hannett, N.M., et al. (2018). Transcription Factors Activate Genes through the Phase-Separation Capacity of Their Activation Domains. Cell 175, 1842–1855.e16.

9. ten Bosch, J.R., Benavides, J.A., and Cline, T.W. (2006). The TAGteam DNA motif controls the timing of Drosophila pre-blastoderm transcription. Development 133, 1967–1977.

10. Bothma, J.P., Garcia, H.G., Esposito, E., Schlissel, G., Gregor, T., and Levine, M. (2014). Dynamic regulation of eve stripe 2 expression reveals transcriptional bursts in living Drosophila embryos. Proc. Natl. Acad. Sci. U. S. A. 111, 10598–10603.

11. Chan, S.H., Tang, Y., Miao, L., Darwich-Codore, H., Vejnar, C.E., Beaudoin, J.D., Musaev, D., Fernandez, J.P., Benitez, M.D.J., Bazzini, A.A., et al. (2019). Brd4 and P300 Confer Transcriptional Competency during Zygotic Genome Activation. Dev. Cell 49, 867–881.e8.

12. Chen, Y.H., Keegan, S., Kahli, M., Tonzi, P., Fenyö, D., Huang, T.T., and Smith, D.J. (2019). Transcription shapes DNA replication initiation and termination in human cells. Nat. Struct. Mol. Biol. 26, 67–77.

13. Cho, W.K., Jayanth, N., English, B.P., Inoue, T., Andrews, J.O., Conway, W., Grimm, J.B., Spille, J.H., Lavis, L.D., Lionnet, T., et al. (2016). RNA Polymerase II cluster dynamics predict mRNA output in living cells. Elife 5, 1–31.

14. Cho, W.K., Spille, J.H., Hecht, M., Lee, C., Li, C., Grube, V., and Cisse, I.I. (2018). Mediator and RNA polymerase II clusters associate in transcription-dependent condensates. Science (80-.). 361, 412–415.

15. Cisse, I.I., Izeddin, I., Causse, S.Z., Boudarene, L., Senecal, A., Muresan, L., Dugast-Darzacq, C., Hajj, B., Dahan, M., and Darzacq, X. (2013). Real-time dynamics of RNA polymerase II clustering in live human cells. Science (80-.). 341, 664–667.

16. Colonnetta, M.M., Abrahante, J.E., Schedl, P., Gohl, D.M., and Deshpande, G. (2021). CLAMP regulates zygotic genome activation in Drosophila embryos. Genetics 219.

17. Comoglio, F., Schlumpf, T., Schmid, V., Rohs, R., Beisel, C., and Paro, R. (2015). High-Resolution Profiling of Drosophila Replication Start Sites Reveals a DNA Shape and Chromatin Signature of Metazoan Origins. Cell Rep. 11, 821–834.

18. Cramer, P. (2019). Organization and regulation of gene transcription. Nature 573, 45–54.

19. Duan, J.E., Rieder, L.E., Colonnetta, M.M., Huang, A., McKenney, M., Watters, S., Deshpande, G., Jordan, W.T., Fawzi, N.L., and Larschan, E.N. (2021). Clamp and zelda function together to promote drosophila zygotic genome activation. Elife 10, e69937.

20. Dufourt, J., Trullo, A., Hunter, J., Fernandez, C., Lazaro, J., Dejean, M., Morales, L., Nait-Amer, S., Schulz, K.N., Harrison, M.M., et al. (2018). Temporal control of gene expression by the pioneer factor Zelda through transient interactions in hubs. Nat. Commun. 9, 1–13.

21. Duronio, R.J., and Marzluff, W.F. (2017). Coordinating cell cycle-regulated histone gene expression through assembly and function of the Histone Locus Body. RNA Biol. 14.

22. Edgar, B.A., and Schubiger, G. (1986). Parameters controlling transcriptional activation during early drosophila development. Cell 44, 871–877.

23. Erickson, J.W., and Cline, T.W. (1993). A bZIP protein, sisterless-a, collaborates with bHLH transcription factors early in Drosophila development to determine sex. Genes Dev. 7, 1688–1702.

24. Espinola, S.M., Götz, M., Bellec, M., Messina, O., Fiche, J.B., Houbron, C., Dejean, M., Reim, I., Cardozo Gizzi, A.M., Lagha, M., et al. (2021). Cis-regulatory chromatin loops arise before TADs and gene activation, and are independent of cell fate during early Drosophila development. Nat. Genet. 53, 477–486.

25. Forero-Quintero, L.S., Raymond, W., Handa, T., Saxton, M.N., Morisaki, T., Kimura, H., Bertrand, E., Munsky, B., and Stasevich, T.J. (2021). Live-cell imaging reveals the spatiotemporal organization of endogenous RNA polymerase II phosphorylation at a single gene. Nat. Commun. 12, 3158.

26. García-Muse, T., and Aguilera, A. (2016). Transcription-replication conflicts: How they occur and how they are resolved. Nat. Rev. Mol. Cell Biol. 17, 553–563.

27. Garcia, H.G., Tikhonov, M., Lin, A., and Gregor, T. (2013). Quantitative Imaging of Transcription in Living Drosophila Embryos Links Polymerase Activity to Patterning. Curr. Biol. 23, 2140–2145.

28. Gaskill, M.M., Gibson, T.J., Larson, E.D., and Harrison, M.M. (2021). GAF is essential for zygotic genome activation and chromatin accessibility in the early Drosophila embryo. Elife 10, e66668.

29. Gratz, S.J., Cummings, A.M., Nguyen, J.N., Hamm, D.C., Donohue, L.K., Harrison, M.M., Wildonger, J., and O’Connor-Giles, K.M. (2013). Genome Engineering of Drosophila with the CRISPR RNA-Guided Cas9 Nuclease. Genetics 194, 1029-+.

30. Hamm, D.C., Larson, E.D., Nevil, M., Marshall, K.E., Bondra, E.R., and Harrison, M.M. (2017). A conserved maternal-specific repressive domain in Zelda revealed by Cas9-mediated mutagenesis in Drosophila melanogaster. PLoS Genet. 13, 1–22.

31. Harlen, K.M., and Churchman, L.S. (2017). The code and beyond: Transcription regulation by the RNA polymerase II carboxy-terminal domain. Nat. Rev. Mol. Cell Biol. 18, 263–273.

32. Harrison, M.M., Li, X.-Y., Kaplan, T., Botchan, M.R., and Eisen, M.B. (2011). Zelda binding in the early Drosophila melanogaster embryo marks regions subsequently activated at the maternal-to-zygotic transition. PLoS Genet. 7, e1002266–13.

33. Henninger, J.E., Oksuz, O., Shrinivas, K., Sagi, I., LeRoy, G., Zheng, M.M., Andrews, J.O., Zamudio, A. V., Lazaris, C., Hannett, N.M., et al. (2021). RNA-Mediated Feedback Control of Transcriptional Condensates. Cell 184, 207–225.e24.

34. Hnisz, D., Shrinivas, K., Young, R.A., Chakraborty, A.K., and Sharp, P.A. (2017). A Phase Separation Model for Transcriptional Control. Cell 169, 13–23.

35. Hsiung, C.C.S., Bartman, C.R., Huang, P., Ginart, P., Stonestrom, A.J., Keller, C.A., Face, C., Jahn, K.S., Evans, P., Sankaranarayanan, L., et al. (2016). A hyperactive transcriptional state marks genome reactivation at the mitosis-G1 transition. Genes Dev. 30, 1423–1439.

36. Huang, S.-K., Whitney, P.H., Dutta, S., Shvartsman, S.Y., and Rushlow, C.A. (2021). Spatial organization of transcribing loci during early genome activation in Drosophila. Curr. Biol.

37. Hur, W., Kemp, J.P., Tarzia, M., Deneke, V.E., Marzluff, W.F., Duronio, R.J., and Di Talia, S. (2020). CDK-Regulated Phase Separation Seeded by Histone Genes Ensures Precise Growth and Function of Histone Locus Bodies. Dev. Cell 54, 379–394.e6.

38. Ji, J.Y., Squirrell, J.M., and Schubiger, G. (2004). Both Cyclin B levels and DNA-replication checkpoint control the early embryonic mitoses in Drosophila. Development 131, 401–411.

39. Kemp, J.P., Yang, X.C., Dominski, Z., Marzluff, W.F., and Duronio, R.J. (2021). Superresolution light microscopy of the Drosophila histone locus body reveals a core-shell organization associated with expression of replication-dependent histone genes. Mol. Biol. Cell 32, 942–955.

40. Kwasnieski, J.C., Orr-Weaver, T.L., and Bartel, D.P. (2019). Early genome activation in Drosophila is extensive with an initial tendency for aborted transcripts and retained introns. Genome Res. 29, 1188–1197.

41. Li, X.-Y., Harrison, M.M., Villalta, J.E., Kaplan, T., and Eisen, M.B. (2014). Establishment of regions of genomic activity during the Drosophila maternal to zygotic transition. Elife 3, e1003428.

42. Liang, H.-L., Nien, C.-Y., Liu, H.-Y., Metzstein, M.M., Kirov, N., and Rushlow, C. (2008). The zinc-finger protein Zelda is a key activator of the early zygotic genome in Drosophila. Nature 456, 400–403.

43. Lucas, T., Ferraro, T., Roelens, B., De Las Heras Chanes, J., Walczak, A.M., Coppey, M., and Dostatni, N. (2013). Live Imaging of Bicoid-Dependent Transcription in Drosophila Embryos. Curr. Biol. 23, 2135–2139.

44. Ludlam, W.H., Taylor, M.H., Tanner, K.G., Denu, J.M., Goodman, R.H., and Smolik, S.M. (2002). The Acetyltransferase Activity of CBP Is Required for wingless Activation and H4 Acetylation in Drosophila melanogaster. Mol. Cell. Biol. 22, 3832–3841.

45. McCleland, M.L., Shermoen, A.W., and O’Farrell, P.H. (2009). DNA replication times the cell cycle and contributes to the mid-blastula transition in Drosophilaembryos. J. Cell Biol. 187, 7–14.

46. McDaniel, S.L., Gibson, T.J., Schulz, K.N., Fernandez Garcia, M., Nevil, M., Jain, S.U., Lewis, P.W., Zaret, K.S., and Harrison, M.M. (2019). Continued Activity of the Pioneer Factor Zelda Is Required to Drive Zygotic Genome Activation. Mol. Cell 74, 185–195.e4.

47. Merrikh, H., Zhang, Y., Grossman, A.D., and Wang, J.D. (2012). Replication-transcription conflicts in bacteria. Nat. Rev. Microbiol. 10, 449–458.

48. Mir, M., Reimer, A., Haines, J.E., Li, X.Y., Stadler, M., Garcia, H., Eisen, M.B., and Darzacq, X. (2017). Dense bicoid hubs accentuate binding along the morphogen gradient. Genes Dev. 31, 1784–1794.

49. Mir, M., Stadler, M.R., Ortiz, S.A., Hannon, C.E., Harrison, M.M., Darzacq, X., and Eisen, M.B. (2018). Dynamic multifactor hubs interact transiently with sites of active transcription in drosophila embryos. Elife 7, 1– 27.

50. Nien, C.-Y., Liang, H.-L., Butcher, S., Sun, Y., Fu, S., Gocha, T., Kirov, N., Manak, J.R., and Rushlow, C. (2011). Temporal Coordination of Gene Networks by Zelda in the Early Drosophila Embryo. PLoS Genet. 7, e1002339–16.

51. Nizami, Z., Deryusheva, S., and Gall, J.G. (2010). The Cajal body and histone locus body. Cold Spring Harb. Perspect. Biol. 2, a000653.

52. Ogiyama, Y., Schuettengruber, B., Papadopoulos, G.L., Chang, J.M., and Cavalli, G. (2018). Polycomb-Dependent Chromatin Looping Contributes to Gene Silencing during Drosophila Development. Mol. Cell 71, 73–88.e5.

53. Palozola, K.C., Donahue, G., Liu, H., Grant, G.R., Becker, J.S., Cote, A., Yu, H., Raj, A., and Zaret, K.S. (2017). Mitotic transcription and waves of gene reactivation during mitotic exit. Science (80-.). 358, 119–122.

54. Park, J., Estrada, J., Johnson, G., Vincent, B.J., Ricci-Tam, C., Bragdon, M.D., Shulgina, Y., Cha, A., Wunderlich, Z., Gunawardena, J., et al. (2019). Dissecting the sharp response of a canonical developmental enhancer reveals multiple sources of cooperativity. Elife 8.

55. Petryk, N., Kahli, M., D’Aubenton-Carafa, Y., Jaszczyszyn, Y., Shen, Y., Silvain, M., Thermes, C., Chen, C.L., and Hyrien, O. (2016). Replication landscape of the human genome. Nat. Commun. 7, 10208.

56. Pourkarimi, E., Bellush, J.M., and Whitehouse, I. (2016). Spatiotemporal coupling and decoupling of gene transcription with DNA replication origins during embryogenesis in C. elegans. Elife 5, 1–12.

57. Rieder, L.E., Koreski, K.P., Boltz, K.A., Kuzu, G., Urban, J.A., Bowman, S.K., Zeidman, A., Jordan, W.T., Tolstorukov, M.Y., Marzluff, W.F., et al. (2017). Histone locus regulation by the Drosophila dosage compensation adaptor protein CLAMP. Genes Dev. 31.

58. Rodríguez-Martínez, M., Pinzón, N., Ghommidh, C., Beyne, E., Seitz, H., Cayrou, C., and Méchali, M. (2017). The gastrula transition reorganizes replication-origin selection in Caenorhabditis elegans. Nat. Struct. Mol. Biol. 24, 290–299.

59. Sabari, B.R., Dall’Agnese, A., Boija, A., Klein, I.A., Coffey, E.L., Shrinivas, K., Abraham, B.J., Hannett, N.M., Zamudio, A. V., Manteiga, J.C., et al. (2018). Coactivator condensation at super-enhancers links phase separation and gene control. Science (80-.). 361, eaar3958.

60. Schindelin, J., Arganda-Carreras, I., Frise, E., Kaynig, V., Longair, M., Pietzsch, T., Preibisch, S., Rueden, C., Saalfeld, S., Schmid, B., et al. (2012). Fiji: An open-source platform for biological-image analysis. Nat. Methods 9, 676–682.

61. Schulz, K.N., and Harrison, M.M. (2019). Mechanisms regulating zygotic genome activation. Nat. Rev. Genet. 20, 221–234.

62. Schulz, K.N., Bondra, E.R., Moshe, A., Villalta, J.E., Lieb, J.D., Kaplan, T., McKay, D.J., and Harrison, M.M. (2015). Zelda is differentially required for chromatin accessibility, transcription factor binding, and gene expression in the early Drosophila embryo. Genome Res. 25, 1715–1726.

63. Seller, C.A., and O’Farrell, P.H. (2018). Rif1 prolongs the embryonic S phase at the Drosophila mid-blastula transition. PLoS Biol. 16, e2005687.

64. Seller, C.A., Cho, C.-Y., and O’Farrell, P.H. (2019). Rapid embryonic cell cycles defer the establishment of heterochromatin by Eggless/SetDB1 in Drosophila. Genes Dev. 33, 403–417.

65. Shermoen, A.W., McCleland, M.L., and O’Farrell, P.H. (2010). Developmental Control of Late Replication and S Phase Length. Curr. Biol. 20, 2067–2077.

66. Sheu, Y.-J., and Stillman, B. (2010). The Dbf4-Cdc7 kinase promotes S phase by alleviating an inhibitory activity in Mcm4. Nature 463, 113–117.

67. Sonneville, R., Craig, G., Labib, K., Gartner, A., and Blow, J.J. (2015). Both Chromosome Decondensation and Condensation Are Dependent on DNA Replication in C.elegans Embryos. Cell Rep. 12, 405–417.

68. Stephenson, R., Hosler, M.R., Gavande, N.S., Ghosh, A.K., and Weake, V.M. (2015). Characterization of a Drosophila Ortholog of the Cdc7 Kinase A ROLE FOR Cdc7 IN ENDOREPLICATION INDEPENDENT OF CHIFFON. J. Biol. Chem. 290, 1332–1347.

69. Strong, I.J.T., Lei, X., Chen, F., Yuan, K., and O’Farrell, P.H. (2020). Interphase-arrested Drosophila embryos activate zygotic gene expression and initiate mid-blastula transition events at a low nuclear-cytoplasmic ratio. PLoS Biol. 18, e3000891.

70. Sun, Y., Nien, C.-Y.Y., Chen, K., Liu, H.-Y.Y., Johnston, J., Zeitlinger, J., and Rushlow, C. (2015). Zelda overcomes the high intrinsic nucleosome barrier at enhancers during Drosophilazygotic genome activation. Genome Res. 25, 1703–1714.

71. Terzo, E.A., Lyons, S.M., Poulton, J.S., Temple, B.R.S., Marzluff, W.F., and Duronio, R.J. (2015). Distinct self-interaction domains promote Multi Sex Combs accumulation in and formation of the Drosophila histone locus body. Mol. Biol. Cell 26.

72. Tie, F., Banerjee, R., Stratton, C.A., Prasad-Sinha, J., Stepanik, V., Zlobin, A., Diaz, M.O., Scacheri, P.C., and Harte, P.J. (2009). CBP-mediated acetylation of histone H3 lysine 27 antagonizes Drosophila Polycomb silencing. Development 136, 3131–3141.

73. White, A.E., Burch, B.D., Yang, X.C., Gasdaska, P.Y., Dominski, Z., Marzluff, W.F., and Duronio, R.J. (2011). Drosophila histone locus bodies form by hierarchical recruitment of components. J. Cell Biol. 193.

74. Yamada, S., Whitney, P.H., Huang, S.K., Eck, E.C., Garcia, H.G., and Rushlow, C.A. (2019). The Drosophila Pioneer Factor Zelda Modulates the Nuclear Microenvironment of a Dorsal Target Enhancer to Potentiate Transcriptional Output. Curr. Biol. 29, 1387–1393.e5.

75. Zaret, K.S., and Carroll, J.S. (2011). Pioneer transcription factors: Establishing competence for gene expression. Genes Dev. 25, 2227–2241.

